# Deconvolving clinically relevant cellular immune crosstalk from bulk gene expression using CODEFACS and LIRICS

**DOI:** 10.1101/2021.01.20.427515

**Authors:** Kun Wang, Sushant Patkar, Joo Sang Lee, E. Michael Gertz, Welles Robinson, Fiorella Schischlik, David R. Crawford, Alejandro A. Schäffer, Eytan Ruppin

## Abstract

The tumor microenvironment (TME) is a complex mixture of cell types whose interactions affect tumor growth and clinical outcome. To discover such interactions, we developed CODEFACS (COnfident DEconvolution For All Cell Subsets), a tool deconvolving cell-type-specific gene expression in each sample from bulk expression, and LIRICS (LIgand Receptor Interactions between Cell Subsets), a statistical framework prioritizing clinically relevant ligand-receptor interactions between cell types from the deconvolved data. We first demonstrate the superiority of CODEFACS versus the state-of-the-art deconvolution method, CIBERSORTx. Second, analyzing the TCGA, we uncover cell-type-specific interactions of mismatch-repair-deficient tumors that are associated with their higher anti-PD1 response rates, including specific T-cell co-stimulating interactions that enhance immunotherapy response independently of the tumors mutation burden levels. Finally, we identify a subset of ligand-receptor interactions in the melanoma TME that predict patient response to anti-PD1 therapy better than recently published transcriptomics-based methods.

## Introduction

The importance of the tumor microenvironment (TME) in cancer has been recognized since the late 1800s^1^. The recent success of immune checkpoint blockade has further sparked interest in studying the various cellular states of cell types and their interactions in the TME, aiming to find biomarkers of treatment response and new treatment opportunities^2^. One key step in elucidating the different cellular states of cell types *de novo* is the characterization of the molecular profiles of each cell type in *each patient’s tumor sample*. Fluorescence-activated cell sorting (FACS) and single-cell RNA sequencing have emerged as effective tools to address this challenge^3^. The use of FACS is limited because it can only measure expression of a few protein markers simultaneously. The use of single-cell RNA sequencing has been limited due to its cost and the scarcity of fresh tumor biopsies. Since bulk tumor gene expression from preserved biopsies accompanied by clinical outcome metadata is abundant, computational methods that can effectively extract cell-type-specific expression from such data in each individual sample could be very helpful. If successful, such deconvolution methods could be used to identify clinically relevant cellular states of cell types and prioritize cell-cell interactions associated with patient survival and response to treatment. Furthermore, they may be readily applied to interrogate the troves of large bulk expression datasets that are in the public domain, to further advance what we can learn from them.

Recovery of cell-type-specific gene expression profiles from bulk gene expression is a computationally challenging problem because the standard formulation requires one to solve a large system of underdetermined linear equations. This problem is closely related to the problem of compressed sensing in signal processing^4^. Several recent studies have developed a variety of algorithms to address this challenge. DeMixT^5^ was designed to estimate individual-specific expression for three cell components provided prior reference samples of two of these cell components. ISOpure^6^ has aimed to derive sample-specific tumor cell expression only, with the assumption that the observed bulk gene expression profile is a mixture of predefined stromal and immune cell expression profiles that are shared across all the samples. Building on this work, Fox et al extended ISOpure to predict individual-specific non-tumor cell expression by subtracting cancer cell expression profiles from the bulk mixtures in a two-cell type model^7^. More recently, Newman et al^8^ developed CIBERSORTx, the first approach that aims to predict the *sample-specific* gene expression of all cell types composing it by employing a set of novel deconvolution heuristics. As a proof of concept, Newman et al^8^ showed that CIBERSORTx can accurately reconstruct the cell-type-specific expression of genes in each input sample under certain modelling assumptions. This groundbreaking work has, however, some notable limitations: (1) The number of genes whose cell-type-specific expression can be reconstructed in each sample is quite small, especially for low-abundance cell types, and (2) CIBERSORTx does not provide confidence estimates of the predictions made. Confidence estimates on the expression predictions could be useful in most deconvolution applications because of the absence of ground truth data.

Starting from the work of Newman et al^8^, we introduce a new deconvolution algorithm and software, CODEFACS (COnfident DEconvolution For All Cell Subsets) that markedly advances our ability to successfully reconstruct cell-type-specific gene expression of each bulk sample. CODEFACS receives as input bulk gene expression profiles of tumor samples and either pre-computed estimates of abundance of expected tumor, immunological and stromal cell types in each sample, or their prototypical molecular signatures, which serve as seeds for estimating the abundance of each cell type in each sample. CODEFACS then predicts the cell type-specific gene expression profiles in each sample. It is a heuristic approach aimed at maximizing the number of genes in each cell type whose expression across the samples can be estimated as confidently predicted. Using 15 benchmark datasets where the ground-truth is known, we show that CODEFACS robustly improves over CIBERSORTx, both in terms of gene coverage and the individual gene expression estimation accuracy. We focus our comparisons on CIBERSORTx because it is widely used (cited more than 350 times as measured by Google Scholar since its publication in 2019) and to our knowledge, it is the only widely used package that makes sample-specific estimates of gene expression for each cell type.

We additionally introduce LIRICS (LIgand Receptor Interactions between Cell Subsets), that integrates the output of CODEFACS with a database of prior immunological knowledge that we curated to infer the cell-type-specific ligand-receptor pairs that are likely to interact in a given sample. These data can then be analyzed in conjunction with any sample-associated clinical annotations (e.g., response to treatment) to systematically prioritize clinically relevant immune related interactions between any pair of cell types in a given patient’s cancer cohort. This opens up the possibility of identifying phenotypic/clinical associations of individual interactions in large clinical tumor expression cohorts.

Building on the enhanced coverage and accuracy of CODEFACS, we applied it to reconstruct the cell-type-specific transcriptomes of ~8000 tumor samples from 21 cancer types in the Cancer Genome Atlas (TCGA). Analyzing these fully deconvolved TCGA expression datasets using LIRICS, we find a shared repertoire of cell-type-specific ligand-receptor pairs that are likely to interact in the TME of mismatch repair deficient tumors of different tissues of origin. These interactions are associated with improved overall patient survival and high response rates to anti-PD1 treatment, independently of their mutation burden levels. Finally, using machine learning techniques, we identify a subset of intercellular TME interactions that are predictive of survival of melanoma patients receiving immune checkpoint blockade therapy better than recently published transcriptomics-based methods.

In summary, CODEFACS and LIRICS effectively build upon the statistical power of large bulk RNA-seq datasets and prior immunological knowledge to characterize sample specific cell-type-specific gene expression and learn more about the association of different tumor-immune interactions with different clinical measures. Notably, the potential scope of applications of both CODEFACS and LIRICS goes beyond studying the TME, as these tools can be applied to study any disease of interest given bulk gene expression data and relevant reference signatures of cell types involved.

## Results

### Overview of CODEFACS and LIRICS

CODEFACS is designed to characterize the tumor microenvironment by reconstructing the cell-type-specific transcriptomes of each sample from bulk expression. It takes as input the bulk RNA-seq expression values of a cohort of tumor samples and either the estimations of the cell fractions of a pre-defined set of cell types in each sample or their cell-type-specific molecular signature profiles, derived based on reference datasets or from the literature^8^.

CODEFACS then employs a heuristic approach that sequentially executes three modules: *(module 1) high resolution deconvolution, (module 2) hierarchical deconvolution and (module 3) imputation*. In module 1, we perform a high-resolution deconvolution, which extends the CIBERSORTx algorithm. In module 2 (hierarchical deconvolution), bulk expression is modeled as a mixture of two components: a specific cell type of interest and all the remaining cell types. The expression for the cell type of interest is predicted by removing the estimated expression in the second component (using high-resolution deconvolution from module 1) from the bulk mixture. In module 3 – imputation-based deconvolution, we impute the cell-type-specific expression of a specific gene based on the predicted cell-type-specific expression of other high-confidence genes that are co-expressed with that gene in the bulk. Each module is designed to overcome the shortcomings of its predecessor based on their respective modeling assumptions. The confidence ranking system is responsible for classifying all the predictions at the end of each module into high-confidence or low-confidence predictions. Genes classified into the low-confidence class at the end of one module (e.g. module 1) are passed to the next module (e.g. module 2) for refinement. Finally, after all three modules are executed, the prediction confidence levels are re-evaluated. The final output of CODEFACS consists of a 3-dimensional matrix with cell-type-specific gene expression predictions for each sample, along with a 2-dimensional matrix of estimated confidence scores of predictions for each gene in each cell type. For more details, see Supplementary Note 1.

A key component of CODEFACS is its *confidence ranking system*, which receives cell-type-specific expression predictions from the different modules and labels them as high or low-confidence estimations. Genes whose expression is determined with high confidence in a given module are added to the output set, while low confidence predictions are continued to be processed in subsequent modules (See **Fig. 1A** and **Supplementary Note 1**). The final output of CODEFACS consists of two items: **(a)** a three-dimensional gene expression matrix, where each entry represents the predicted gene expression a gene in a given cell-type in a specific sample, and **(b)** a two-dimensional matrix of confidence scores ranging from [0,1] representing which gene-cell-type pairs have the most confident predictions (**Fig. 1A, Output**).

**Figure 1.**
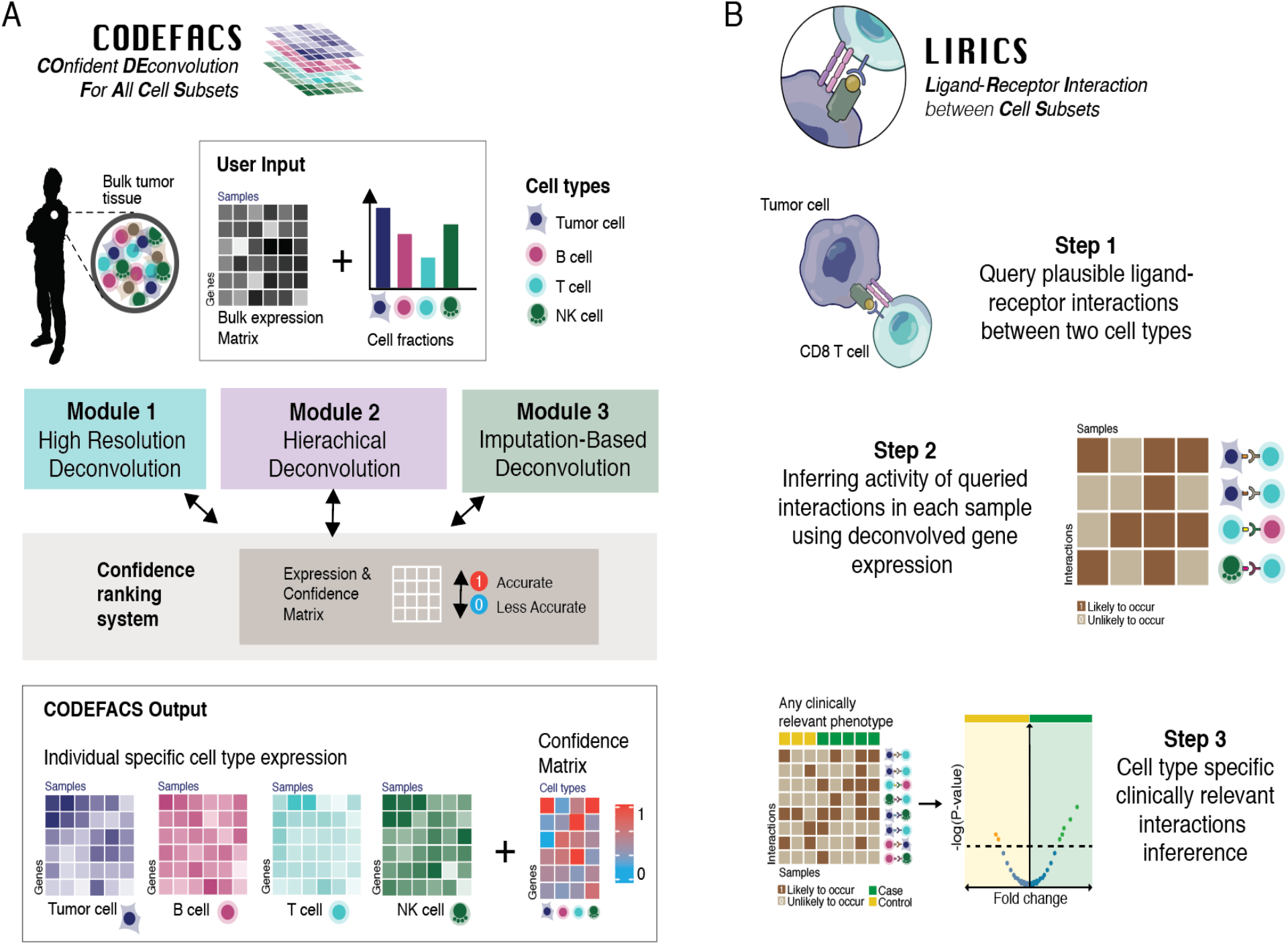
Overview of CODEFACS and LIRICS. **(A) CODEFACS** takes bulk gene expression profiles and prior knowledge of the cellular composition of each sample and executes a heuristic three-step procedure to infer the deconvolved gene expression in each sample, as described in above and in more details in **Supplementary Note 1**. **(B) LIRICS** takes the output of CODEFACS and processes it in three steps, as described in the main text and in detail in details, see **Supplementary Note 2**.

Given fully deconvolved gene expression data from CODEFACS, one can use LIRICS (LIgand Receptor Interactions between Cell Subsets) (**Fig. 1B**) to infer which cell-type specific ligand-receptor pairs are likely to interact in each sample. Specifically, LIRICS takes the output of CODEFACS and processes it in three steps: (*Step 1*) The first step queries a database of all plausible ligand-receptor interactions between any two cell types A and B, which we have systematically assembled and curated from the literature. This database is publicly available as part of LIRICS (see **Supplementary Note 2**, **Supplementary Tables 1-3**) and serves as a repository of all prior immunological knowledge. (*Step 2*) In the second step, given the deconvolved expression profiles of cell type A and cell type B in a given bulk tumor sample, LIRICS denotes as ‘active’ or ‘likely to occur’ (‘1’) the interactions where both the ligand and receptor are over-expressed in the relevant cell-types in that sample, or otherwise ‘inactive’ (‘0’). A ligand or receptor is considered to be overexpressed in a given cell type if its expression exceeds the median expression in that cell type (**Supplementary Note 2**). (*Step 3*) In the third step, a Fisher’s enrichment analysis is performed to test the association of the activity of specific ligand-receptor interactions with any relevant phenotypes of interes (e.g., treatment response, mutational subtype, etc.) (**Fig. 1B**).

### Evaluating the performance of CODEFACS versus numerous different benchmarks

To assess the accuracy of CODEFACS, we generated 15 benchmark datasets (see Methods) by merging publicly available single cell RNA-seq^9,10^ and FACS sorted purified RNA-seq^11^. Thereafter, we applied CODEFACS to deconvolve these generated bulk datasets and define the accuracy of its predictions by computing the Kendall correlation between the predicted and ground truth expression in each cell type across individual samples (the Kendall correlation provides a less inflated measure of accuracy by accounting for ties in the data). In the main text, we show the results obtained on three of these benchmark bulk datasets: one derived form a FACS-sorted lung cancer data, one from a single cell melanoma RNA-seq data and from a single cell glioblastoma RNA-seq dataset, respectively. Each bulk sample from these datasets represents a biopsy from a patient. We show that CODEFACS predicts the cell-type-specific expression of many more genes than CIBERSORTx (with Kendal’s correlation ≥ 0.3) (**Fig. 2A-C**), and its predictions are overall more accurate (**Fig. 2D-F**). The results for all the remaining 12 benchmark datasets, created via artificial mixing of single cell profiles and simulation of batch effects between bulk and single cell expression data (See Methods and **Supplementary Table 4** for more details), further show the superiority of CODEFACS (**Supplementary Fig. S1 and S2**). Overall, we observe that the more abundant the cell type is, the better CODEFACS can predict its cell-type-specific gene expression (**Supplementary Fig. S3**).

**Figure 2.**
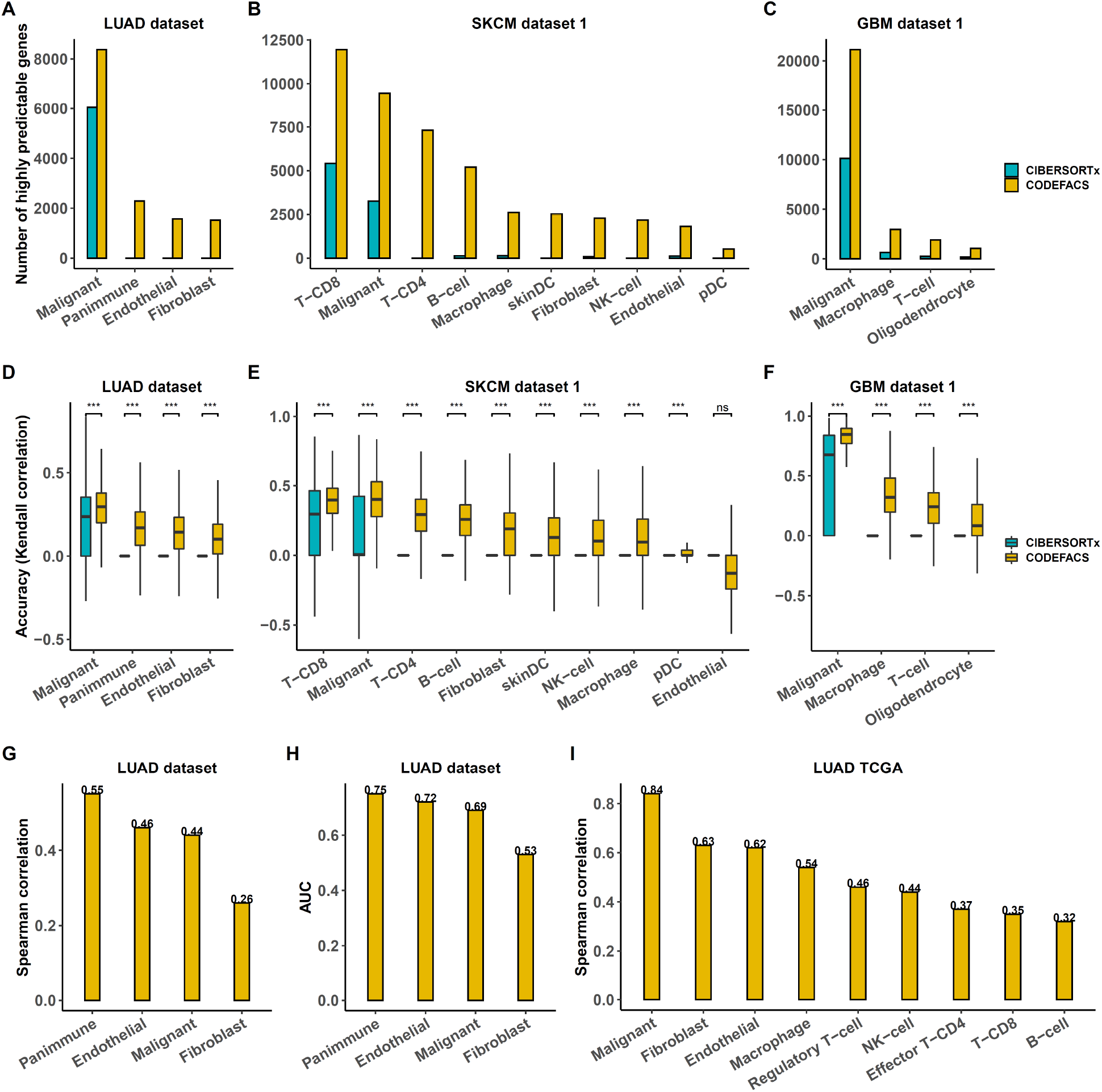
Evaluating the performance of CODEFACS. **(A-C)** bar plots depicting the number of genes with a prediction accuracy (Kendall correlation) ≥ 0.3 with the ground truth for each cell type, estimated from bulk-generated samples of lung cancer (LUAD dataset; sample size = 26), melanoma (SKCM dataset 1; sample size = 28) and glioblastoma (GBM dataset 1; sample size = 24) benchmark datasets, as estimated by CODEFACS (yellow bars) and CIBERSORTx (blue bars). **(D-F)** boxplots depicting prediction accuracy distributions of all genes across different cell types in the lung cancer (LUAD with sample size 26), melanoma (SKCM with sample size 28) and glioblastoma (GBM with sample size 24) benchmark datasets, using CODEFACS (yellow) and CIBERSORTx (blue). A two-sided Wilcoxon signed rank test was performed to compare the prediction accuracies of CODEFACS and that of CIBERSORTx for each cell type in each dataset. *** denotes p-values < 2e-16. **(G)** Spearman correlations between prediction accuracies and confidence scores among cell types in the lung cancer benchmark dataset (LUAD dataset; sample size = 26). The y-axis indicates the spearman correlation coefficient value, while the x-axis indicates the cell type. **(H)** AUCs obtained in classifying informative and uninformative predictions among cell types in lung cancer benchmark dataset (LUAD dataset; sample size = 26). **(I)** bar plots depicting the Spearman correlations between mean deconvolved cell-type-specific expression in TCGA-LUAD and mean cell-type-specific expression derived from publicly available single cell datasets of LUAD. The y-axis indicates the Spearman correlation coefficient value, while the x-axis indicates the cell type. Details of all single cell RNA-Seq based pseudo bulk datasets can be found in **Supplementary Table 4.**

To test CODEFACS confidence scores can filter out potentially noisy predictions we quantified how well do these scores align with the Kendall scores that measure the prediction accuracy with the ground truth for each (gene, cell type) pair, as described above. We quantified this using two metrics: Spearman correlation and a classification Area Under the ROC Curve (AUC). As evident, this correlation is strong and positive (**Fig. 2G** depicts the results for the FACS sorted lung cancer benchmark dataset, and **Supplementary Fig. S4** depicts the results for the remaining benchmark datasets). To perform a classification-based quantification, we grouped the genes in each cell type into two classes based on the correlation between their predicted and actual expression, as *informative* (prediction accuracy ≥ 0.1 and p-value ≤ 0.05) and *uninformative* (prediction accuracy < 0.1 or p-value > 0.05). We then tested whether the confidence scores could be used to classify genes into these two classes for each cell type. We found that the confidence score could effectively filter out uninformative predictions (**Fig. 2H** depicts the results for the FACS sorted lung cancer benchmark dataset, and **Supplementary Fig. S5**, depicts for the remaining benchmark datasets).

Finally, to evaluate CODEFACS on real bulk tumor data, we applied it to deconvolve bulk expression data from 21 cancer types (~8000 RNA-seq samples) in TCGA. To infer the cellular abundance of each cell type in each sample which is required as input for CODEFACS, we made use of matched bulk methylation data available for these samples and methylation-based reference signature profiles of distinct cell types. These include 11 cell type signatures (macrophages/dendritic cells--CD14+, B cells--CD19+, CD4+T cells, CD8+ T cells, T regulatory cells, NK cells--CD56+, endothelial cells, fibroblasts, neutrophils, basophils, eosinophils and tissue-specific tumor cells) obtained from MethylCIBERSORT^12^. Reassuringly, we found high Spearman correlations between the resulting predicted tumor cell fraction and the tumor purity estimates derived from matched mutation and copy number data (based on ABSOLUTE) for the same samples in each of 10 different cancer types (Spearman correlation: min=0.72, max=0.88, avg=0.8; **Supplementary Fig. S6**).

We then asked if CODEFACS can recover the cell-type-specific gene expression signature of different cell types in a given cancer type? To this end, we computed the Spearman correlation between (a) the mean deconvolved gene expression of the top confidently deconvolved genes in a given cell type (confidence score ≥ 0.95) and (b) the mean expression of these genes, which we derived from completely independent single cell expression data of the same cancer type (Methods). We find that (a) and (b) are substantially correlated (**Fig. 2I** depicts results for the TCGA-LUAD (lung adenocarcinoma) dataset as an example and **Supplementary Fig. S7** for the remaining cancer types that have publicly available scRNA-seq data). The concordance level is higher for cell types that are abundant (e.g., tumor cells and fibroblasts) and decreases for less abundant cell types. Additionally, we observed that tumor cells have the largest fraction of genes whose expression is predicted with high confidence, with the highest in thyroid cancer (THCA, ~67.4% of all genes). Furthermore, seven KEGG pathways are significantly enriched (adjusted p-value < 0.01) with highly confident genes in tumor cells (confidence score ≥0.95) across the 21 cancer types (**Supplementary Fig. S8**). Those pathways mostly involve RNA transport, spliceosome, DNA replication, and mismatch repair.

### Tumors with DNA mismatch repair deficiency have heightened T cell co-stimulation that is independent of their tumor mutation burden levels

In normal cells, DNA is constantly repaired in response to DNA damage or DNA replication errors^13^. However, defects in specific DNA repair pathways in cancer cells may result in the accumulation of many somatic mutations resulting in hypermutated tumors (TMB ≥ 10-20 mutation/Mb)^14–16^. One cause of hypermutability is a mismatch repair deficiency (MMRD), which leads to the accumulation of insertions and deletion mutations in microsatellite regions of the genome due to uncorrected DNA replication polymerase slippage events. This is known as microsatellite instability (MSI)^17,18^. Solid tumors with mismatch repair deficiency were shown to be sensitive to immune checkpoint blockade (ICB) therapy irrespective of tumor type, leading the FDA to approve MSI as the first cancer type agnostic biomarker for patients receiving anti-PD1 treatment^19^. The reason behind this general sensitivity to anti-PD1 treatment is not completely understood. Prior work has led to the prevailing hypothesis that elevated tumor mutation burden in mismatch repair deficient tumors leads to more neoantigens, and thus is more likely to activate a host immune response against tumor cells^17,20–22^. However, not all tumor types with elevated tumor mutation burden have similar response rates to anti-PD1^23,24^, and recent studies have revealed that T cells recognize and respond to only a few neoantigens per tumor^25–28^. In the TCGA collection, there are three solid tumor types with a significant association between hypermutability and survival benefit in patients. These three are the MSI enriched solid tumor types (**Fig 3A,B**, borrowed from^29,30^, MSI tumors highlighted as red dots in Fig 3A, **Fig. 3C.** log-rank test p-value = 0.00084). The same cannot be said about other solid tumor types (**Fig 3D**, log-rank test p-value = 0.4). Taken together, these findings motivated us to further study cellular immune crosstalk in the tumor microenvironment of mismatch repair deficient solid tumors to gain additional cell-type-specific insights into their sensitivity to anti-PD1 treatment.

**Figure 3.**
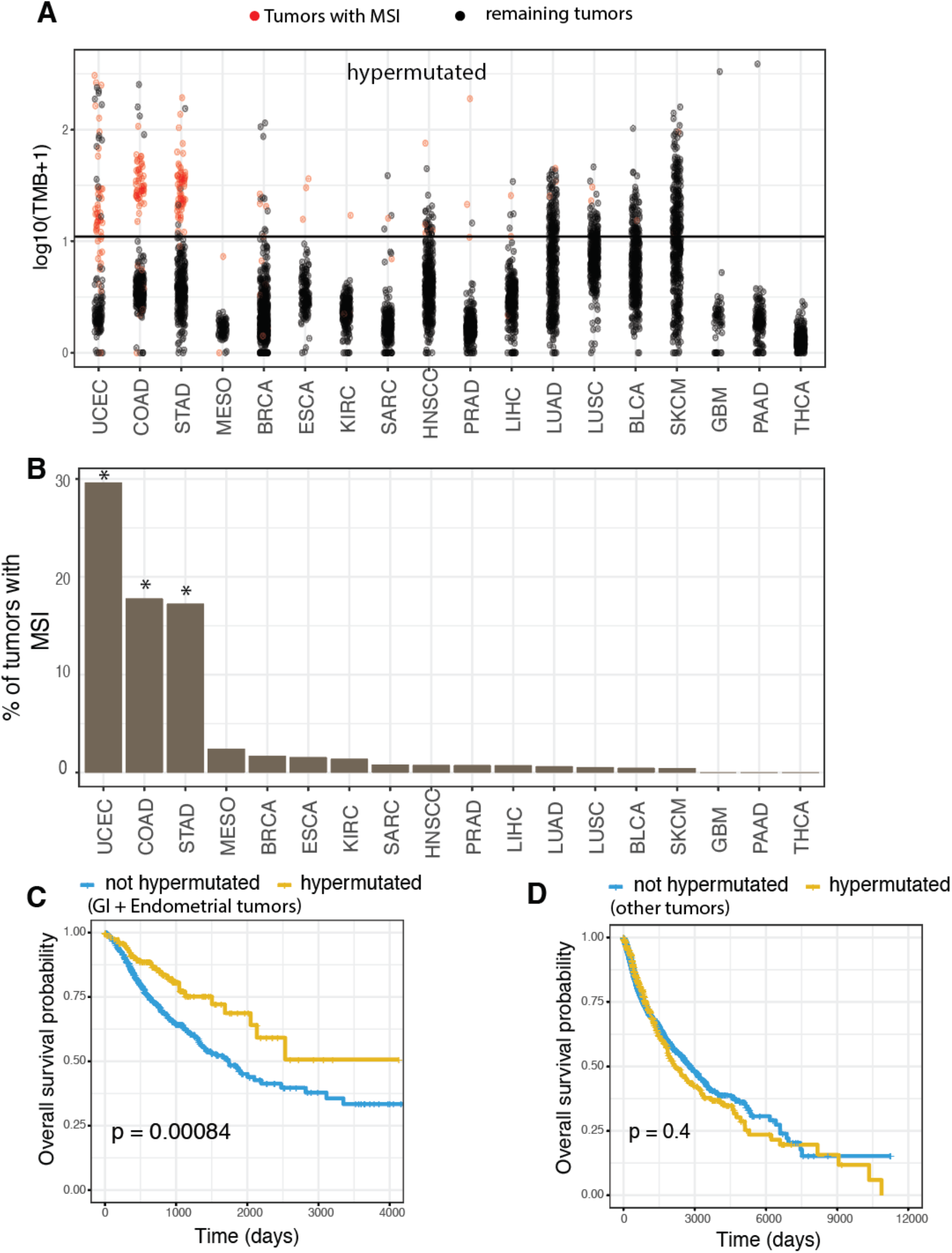
**(A) The landscape of tumor mutation burden and microsatellite instability across 18 different solid tumor types.** This panel plots the distribution of non-synonymous tumor mutation burden on a logarithmic scale (Y-axis). All points above the horizontal line are typically regarded as hyper-mutated tumors (> 10 mutations/Mb). All red points represent tumors with a DNA mismatch repair deficiency detected via microsatellite instability (MSI). **(B) This panel depicts the percentage of all tumor samples per cancer type with microsatellite instability (Y-axis).** Tumor types marked with a * represent those where MSI is prevalent. **(C-D) Comparison of overall survival of patients with tumors that are hypermutated vs not hypermutated.** (**C**) Survival comparison in Gastro-Intestinal+Endometrial tumor types where MSI is prevalent (marked with a * in panel B). (**D**) Survival comparison in all other solid tumor types where tumors rarely have an underlying mismatch repair deficiency. Statistical significance of differences in survival was calculated using the log-rank test.

To identify cell-cell interactions that are differentially active between microsatellite instable tumors and microsatellite stable tumors (red vs black dots in Fig 3A), we applied CODEFACS to deconvolve the bulk gene expression of all solid tumors from TCGA and integrated their predicted cell-type-specific gene expression levels with LIRICS. The top 50 interactions from this differential analysis (ordered by FDR adjusted p-value) are shown in a network in **Fig. 4A**. These interactions are frequently active in hypermutated solid tumors with DNA mismatch repair deficiency compared to other hypermutated tumors (**Fig. 4B, Supplementary Fig. S9**), testifying to their MSI specificity. They include the well established PDL1-PD1 checkpoint interaction between tumor cells and CD8+ T cells, but additionally, numerous T-cell activating/co-stimulatory interactions such as the 41BBL-41BB interaction between tumor cells and CD8+ T cells, ULBP2-NKG2D between tumor cells and CD4+ T cells. They also encompass several chemotaxis related interactions involved in trafficking of lymphocytes in and around the tumor mass, such as the CXCL9-CXCR3 chemokine interaction between macrophages and CD4+ T cells and CCL3/4/5 – CCR5 interactions between various immune and stromal cell-types.

**Figure 4.**
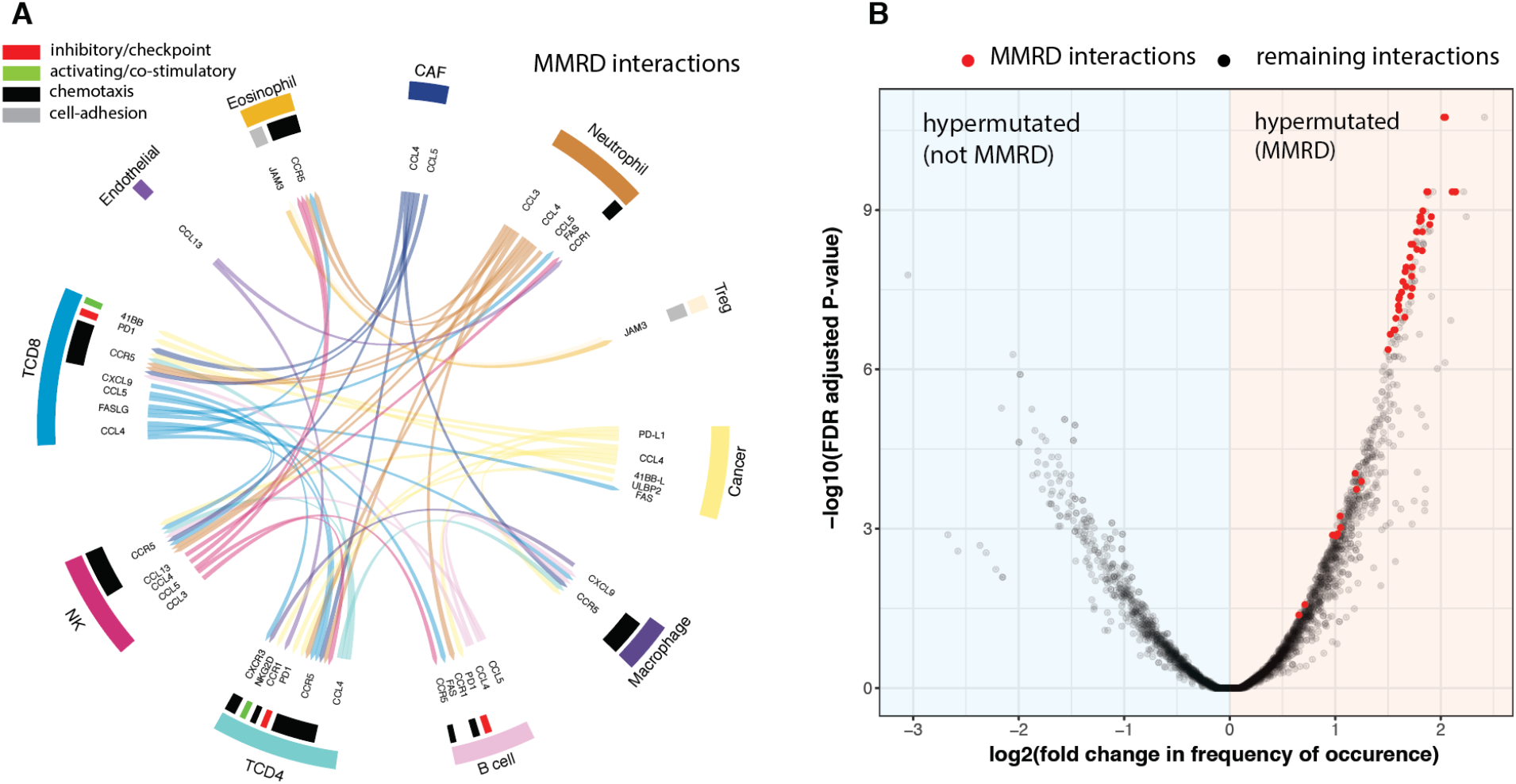
**(A) Interaction network consisting of the top 50 interactions most highly activated in TME of tumors with DNA mismatch repair deficiency.** Interactions highlighted in green represent co-stimulatory interactions/having an activating effect on the target cell. Interactions highlighted in red represent checkpoint interactions/having an inhibitory effect on the target cell. Interactions highlighted in black represent pro-inflammatory/chemotaxis interactions involved in inflammatory response and immune cell trafficking to tumor sites. Eos: Eosinophils, CAF: Cancer associated fibroblasts. **(B) A volcano plot depicting on the x-axis the log2 fold change in the frequency of occurrence of each cell-cell interaction in the TME of hypermutated tumors with an underlying DNA mismatch repair deficiency vs other hypermutated tumors.** The y-axis indicates the −log10 FDR adjusted p-value of the observed enrichment. Highlighted in red in the scatter plot are the top 50 interactions that are most differentially active between all MSI vs non-MSI tumors (shown in panel A).

### Machine learning guided discovery of cell-type-specific ligand-receptor interactions that predict patient response to ICB therapy

We set to discover cell-type-specific ligand-receptor interactions predictive of response to ICB therapy by analyzing the TCGA deconvolved expression we have generated. We focus on melanoma, currently the tumor type best responding to ICB, where there are many independent publicly available bulk expression datasets of patient’s receiving anti-PD1 treatment. Starting from the deconvolved TCGA-SKCM dataset as our training set (N=469), we employed a genetic algorithm to find cell-type specific ligand-receptor interactions, whose activation state best separates hypermutated melanoma tumors from non-hypermutated tumors (see Methods, **Fig 5A**). We term the interactions identified in this process *melanoma hypermutation-specific functional interactions* (*MSFI*), and the network formed by these interactions is displayed in **Fig. 5B**.

**Figure 5.**
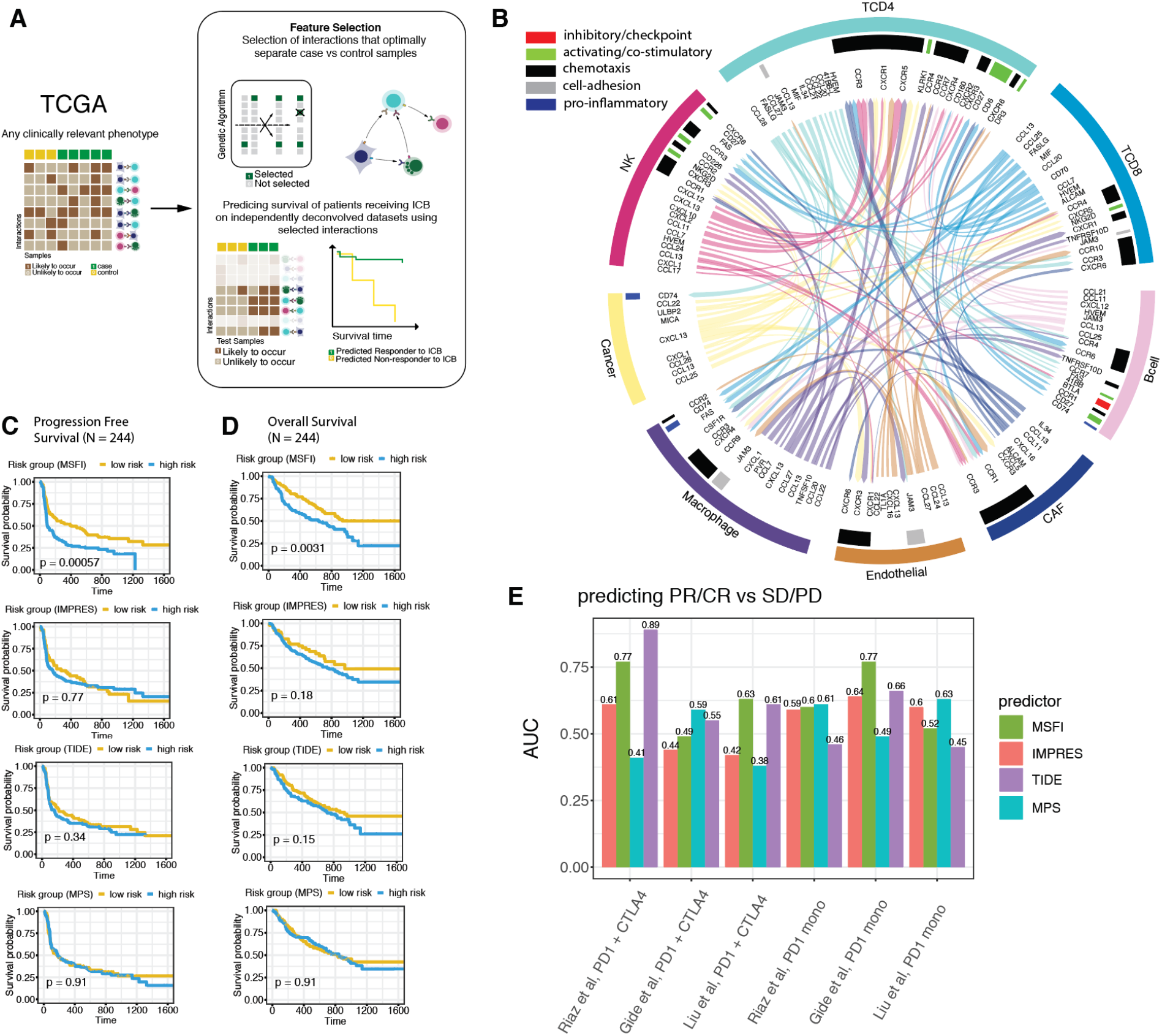
**(A) Overview of the machine learning analysis employed to identify cell type specific interactions that are predictive of response to immune checkpoint blockade therapy. (B) A chord diagram of the resulting MSFI network.** Each individual interaction is represented by a link from the source cell type (ligand expressing cell type) to the target cell type (receptor expressing cell type) and the color of the link represents the color of the source cell type. For interactions that are activating/co-stimulatory, the sector in the corresponding target cell type is highlighted in green. For inhibitory/checkpoint interactions, the sector in the target cell type is highlighted in red. Interactions involved in chemotaxis are highlighted in black and those mediating a pro-inflammatory response are highlighted in blue, cell-adhesion interactions are highlighted in grey. **(C) Kaplan-Meier plot depicting the progression free survival of the combined set of melanoma patients receiving immune checkpoint blockade (N= 244).** On the top, the patients are stratified into low-risk/high-risk groups based on the median value of MSFI score from LIRICS. Second from top, patients stratified into low/high risk groups based on median IMPRES score^46^. Third from top, patients stratified into low/high risk groups based on median TIDE score^47^, Bottom, patients stratified into low/high risk groups based on median MPS score^48^. **(D) Kaplan-Meier plots depicting the overall survival of all melanoma patients receiving immune checkpoint blockade (N= 244).** On the top, the patients are stratified into low-risk/high-risk groups based on the median value of MSFI score from LIRICS. Second from top, patients stratified into low/high risk groups based on median IMPRES score^46^. Third from top, patients stratified into low/high risk groups based on median TIDE score^47^, Bottom, patients stratified into low/high risk groups based on median MPS score^48^. Survival differences among patients that received anti-PD1 monotherapy vs anti-CTLA4 + anti-PD1 combination are shown in **Supplementary Fig. S10 (E) Area under the ROC curves in predicting Complete/Partial-response (based on RECIST v1.1) to immune checkpoint blockade therapy for the different scores.** X-axis marks patients grouped by dataset source and treatment regimen. PD1 mono represents patients that received anti-PD1 monotherapy. PD1 + CTLA4 represents patients that additionally received anti-CTLA4 besides anti-PD1.

Having identified the MSFI interactions, we applied CODEFACS to deconvolve the bulk expression data of pre-treatment samples from the three largest publicly available melanoma datasets where patients received anti-PD1 treatment (either monotherapy or in combination with anti-CTLA4; Methods)^43–45^. We then employed LIRICS (step 1 and step 2) to the respective deconvolved expression of each of these checkpoint datasets, *without any additional training*, and simply quantified the number of MSFI interactions that are active in each of these patients’ tumor samples, which we denote as the tumor’s *MSFI score*. Remarkably, we find that the MSFI score of each sample can robustly stratify patients into those that are likely to respond to ICB vs those that are unlikely to respond (**Fig. 5C**, progression free survival log rank test p-value: 0.00057, **Fig. 5D**, overall survival log rank test p-value: 0.0031). **Supplementary Fig. S10** depicts the survival differences for the two treatment groups separately (anti-PD1 monotherapy and anti-CTLA4 + anti-PD1 combination). Additionally, **Supplementary Fig. S11** and **S12** depict the survival differences for each ICB dataset separately. As evident, our results improve over recent bulk gene expression based predictors of melanoma ICB therapy response (IMPRES^46^, TIDE^47^ and the melanocytic plasticity signature (MPS) scores^48^). We note that the performance levels of the latter on the more recent larger bulk expression datasets, where their original RNAseq reads have been uniformly aligned and normalized as described in the Methods, is lower than that reported in the original publications, pointing to the growing awareness of the sensitivity of expression-based predictors to batch effects, the normalization and alignment methods used, highlighting the need to pre-process the raw sequencing data in a uniform manner in making meaningful comparisons (see Discussion).

To further evaluate the predictive performance of the MSFI score, we tested its ability to predict partial or complete responders vs stable or progressive disease patients in these datasets. To this end, we plotted the receiver operator area under the curve (AUC) obtained using the MSFI score for classifying the patients to partial or complete responders vs stable or progressive disease, and compared its performance to that obtained with the three other published predictors for the different treatment groups (**Fig. 5E**). On average, the MSFI score achieves an AUC of 0.63 (for anti-PD1 monotherapy the AUCs obtained are 0.6, 0.77 and 0.52 for the three individual ICB datasets, and for the anti-CTLA4 + anti-PD1 combination ICB treatment, the AUCs are 0.77, 0.49 and 0.63). A similar performance could not be achieved if the placement of the ligand and receptor between interacting cell-types in the MSFI network was swapped (average AUC 0.58) or by randomly shuffling the interaction activity profiles (average AUC ~ 0.52), testifying that the selected cell-cell interactions are best predictive of response to ICB. Comparing MSFI predictive performance in this response classification task to that of the recent melanoma bulk expression-based predictors, TIDE achieves an average AUC of 0.6 (for anti-PD1 monotherapy the AUCs are 0.46, 0.66 and 0.45 for the three ICB datasets and 0.89, 0.55 and 0.61 for the combination), IMPRES achieves an average AUC of 0.55 (for anti-PD1 monotherapy, the AUCs are 0.59, 0.64 and 0.6 respectively for the three ICB datasets and 0.61, 0.44 and 0.42 for the combination) and MPS achieves an average AUC of 0.51 (for anti-PD1 monotherapy, the AUCs are 0.61, 0.49 and 0.63 respectively for the three ICB datasets and 0.41, 0.59 and 0.38 for the combination).

Examining the MSFI network **(Fig. 5B)**, we find an over-representation of cell-type-specific co-stimulatory/immune cell activating interactions known from prior immunological literature (hypergeometric test p-value < 0.05)^49–59^ (see **Supplementary Table 1**), again highlighting the importance of co-stimulation in mediating successful anti-tumor immune responses. Additionally, the MSFI network additionally includes cytokine/chemokine interactions involved in pro-inflammatory response and the trafficking of NK, T and B cells to the tumor site (responsible for better lymphocyte infiltration into the tumor mass). Importantly, a multivariate Cox-proportional hazards model with MSFI scores and TMB of each patient receiving anti-PD1 (wherever TMB data were available) shows that the MSFI score remains significantly associated with improved progression free survival (p-value: 0.013) and overall survival (p-value: 0.0258) despite differences in mutation burden (TMB PFS p-value: 0.224, OS p-value: 0.477).

Taken together, these results suggest that while recognition of melanoma tumor neo-antigens serves as an initiating trigger, the activation of additional, specific, co-stimulatory and pro-inflammatory signals in the TME could further enhance an effective host immune response upon immune checkpoint blockade treatments.

## Discussion

This study presents a new computational tool, CODEFACS and a supporting immune interactions framework, LIRICS, that enable an (averaged) ‘virtual single cell’ characterization of the TME from bulk tumor expression data. Applying these tools across the TCGA, we systematically identify, for the first time, cell-type-specific ligand-receptor pairs that are likely to interact directly in the TME of specific patient populations and prioritize those that are associated with patients’ response to ICB. Specifically, we identified a shared core of intercellular TME interactions in DNA mismatch repair deficient tumors, which are associated with improved patient survival and high sensitivity to immune checkpoint blockade therapy. Finally, focusing on melanoma, we show that one can bootstrap on the large deconvolved data resource from TCGA using machine learning techniques to discover cell-cell interactions within the TME that successfully predict patients’ response to immune checkpoint blockade.

The heightened cellular crosstalk unique to the TME of mismatch repair deficient tumors suggests that T cells are being activated by co-stimulatory signals in addition to immunogenic neo-antigens only to be kept in balance by other immunoregulatory mechanisms such as the PD1-PDL1 checkpoint interaction between CD8+ T cells and tumor cells. Our results suggest that when this interaction is blocked by anti-PD1 treatment, additional co-stimulatory interactions (such as the 41BBL-41BB between tumor and T cells) lead to the observed enhanced response of MMRD tumors to immune checkpoint blockade therapy. These results further support the possibility that switching on specific T cell co-stimulation signals in the TME may lead to better responses to anti-PD1 treatment independent of tumor mutation burden. Notably, recent pre-clinical studies have shown that combination therapies aimed at enhancing such T-cell co-stimulating interactions improve anti-tumor immune responses even in low TMB and highly immuno-suppressive settings^33–38^. Currently, several clinical trials to assess the safety and efficacy of these combinations are in progress^39–42^.

While we have provided a toolkit to prioritize clinically relevant cell-cell interactions from bulk tumor expression, there are limitations that should be noted and potentially further improved upon in the future. First, several recent publications have reported discrepancies between different RNA-seq expression quantification methods based on the reference transcriptome version used and choice of method (alignment based vs alignment free)^60–64^. This can potentially affect reproducibility of findings. Hence, whenever possible, we recommend that all bulk RNA-seq datasets are homogeneously pre-processed using the same RNA-seq quantification method and reference transcriptome before the application of our tools. Indeed, following this notion, in this study we have pre-processed all bulk RNA-seq datasets in a uniform manner, using STAR (v2.7.6a) + RSEM (v1.3.3) and GENCODE v23 human genome annotation (Methods).

Second, CODEFACS itself has several limitations: (1) it requires prior information about the cell type composition of the input tumors, or alternatively, knowledge of the pertaining cell-types’ gene expression or methylation signatures that can be used to infer their abundances, and its accuracy depends on the accuracy of the latter. (2) It is predictive typically for subsets of the whole exome and its performance deteriorates for lowly-abundant cell types. However, the confidence scores provided help by allowing the user to rank genes in each cell-type by the quality of predictions. Third, LIRICS is currently restricted to well-defined ligand-receptor interactions between tumor, immune, stromal and epithelial cell types and does not consider the spatial localization of cells in the TME. The inclusion of the latter with the advent of forthcoming spatial transcriptomics data is likely to lead to considerably more informative interaction inference approaches.

Although this work focuses on studying the tumor microenvironment, the tools presented here can be applied to prioritize important cell-cell interactions in noncancerous tissues under a variety of normal and disease states. One interesting application that we envision is the characterization of clinically relevant intercellular interactions occurring at the maternal-fetal interface using corresponding bulk gene expression data and pregnancy outcome information, whose elucidation may help treat and mitigate preeclampsia and other pregnancy related complications. One can also use our tools to study bulk gene expression data from pre-malignant tissue samples and compare them against malignant samples to elucidate cell-cell interaction dynamics on the way to malignancy. Finally, one can deconvolve expression data from autoimmune disorders to learn more about the underlying immune interactions. In sum, the computational tools developed and presented here offer a cost-effective way to study immune responses at a cell-type-specific resolution from the widely abundant bulk gene expression in both health and disease, complementing first-line single cell technologies in situations where the latter are less applicable.

## Methods

### Data collection and pre-processing

#### Single cell RNA-seq datasets

To benchmark the performance of CODEFACS, we first set out to obtain publicly available single cell RNA-seq datasets where both tumor and non-tumor cells were successfully isolated. This search led us to the identification of nine such single cell RNA-seq datasets from the literature, each from a different cancer type. Collection of additional single cell datasets was frozen after Dec 2019. Details of the single cell datasets that we curated for this study in **Supplementary Table 5**. For each dataset sequenced on the SmartSeq2 platform, the log normalized transcript counts for each gene in each sequenced cell were made publicly available by the original authors. For the application of deconvolution, these counts were transformed back to the Transcripts Per Million (TPM) scale. For datasets sequenced on the 10x platform, UMI counts for each gene were made publicly available and were scaled by the library size of each cell and multiplied by a factor of 1 million to get expression values in TPM scale.

#### Bulk RNA-seq datasets

Gene expression and matching bulk tumor methylation data from fresh frozen tumor biopsies in TCGA were downloaded from (http://xena.ucsc.edu/)^65^. In addition, publicly available bulk expression data from formalin fixed paraffin embedded tumor biopsies of melanoma patients receiving immune checkpoint blockade treatment were downloaded from^43–45^. All bulk RNA-seq datasets were collected such that they have a sufficiently large sample size to reliably perform complete deconvolution of expression profiles (> 4 times the number of cell-types involved)^8^. Collection of datasets was frozen after Dec 2019. Details on bulk RNA-seq datasets deconvolved in this study can be found in **Supplementary Table 6**. To maintain consistency with the pipeline used for preprocessing TCGA data, bulk gene expression levels in immune checkpoint blockade datasets were re-quantified using STAR v2.7.6a and RSEM v1.3.3^66^ with GENCODE v23 human genome annotation^67^. Furthermore, to mitigate technical biases, between-sample scaling factors were estimated using the TMM method implemented in edgeR^68^ and TPM values in each sample were further rescaled by these scaling factors^69^.

#### Generation of simulated bulk RNA-seq datasets

To evaluate the performance of CODEFACS, we generated 14 different pseudo-bulk RNA-seq datasets from mixing experiments with single-cell data. Each sample in each benchmark dataset has matching cell type specific gene expression profiles derived from averaging single-cell RNA-seq profiles of individual cells from the same sample and same cell type. These profiles serve as the ground truth for the evaluation of deconvolution performance. To avoid any circularity in our validations, for each of the single-cell datasets involved, single cell data from 4 randomly chosen patients were separated from the rest. These data were used to derive reference gene expression signatures for each cell type. The mixing experiments were then performed on single-cell data of the remaining patients that were hidden from the reference signature derivation process. In addition, we simulated technical replicates for each pseudo-bulk sample, wherein we injected noise in the pseudo-bulk expression of a few randomly chosen genes and then renormalized the expression data by the sample library size. This procedure simulates mRNA composition noise that is commonly observed in bulk RNA-seq datasets due to technical differences in sample preparation [64-66]. The datasets and the mixing experiment used to generate the pseudo-bulk samples are listed in **Supplementary Table 4**. In addition, we obtained a FACS sorted lung cancer dataset which include purified RNA-seq for four cell types^11^ and generated a pseudo bulk correspondingly. In total, 15 benchmark datasets were generated.

### Deconvolving immune checkpoint blockade (ICB) and TCGA bulk RNA-seq datasets using CODEFACS

For the application of CODEFACS, molecular profiles of signature genes of each cell type of interest are needed to estimate the relative cell fractions in the bulk. We used single-cell-expression-derived signatures as priors to deconvolve the melanoma ICB datasets. To derive these signatures from single-cell data, we first start out by obtaining the class labels of each cell type of interest. These data are publicly available for each single-cell dataset we collected. Hence, we primarily use these labels in our study (unless further refinement of labels into specific cell subtypes of interest is needed for a specific usage). With a collection of single cell expression profiles and matching cell type labels as input, we used CIBERSORT online tool to derive a cell-type-specific signature matrix. Thereafter, we applied CODEFACS to ICB datasets with default parameters settings and batch correction requirement specified. For TCGA deconvolution, we first estimated cell fractions based on bulk methylation and then applied CODEFACS to corresponding bulk gene expression for the 21 cancer types which have both types of data available. We chose methylation signatures over expression-based signatures for TCGA analysis for two reasons. First, single cell expression data with consistent cell types across 21 cancer types are not available. Second, DNA methylation-based signatures are considered to be more stable marks of cellular identity compared to dynamic RNA expression derived signatures^70^. The methylation-based cell type signatures (**Supplementary Table 7**) were obtained from MethylCIBERSORT^12^. We applied CODEFACS to TCGA datasets with default parameters settings and without batch correction option specified. For the details of CODEFACS algorithm and parameter settings, see **Supplementary Note 1**.

### Machine learning for discovery of cell-cell interactions predictive of clinical responses to immune checkpoint blockade in melanoma

We used a genetic algorithm, which is a randomized heuristic search algorithm designed to select optimal features for a prediction task given some user-defined fitness function for training^71^. In this setting, the features are ligand-receptor interactions between cell-types, the prediction task is predicting response to ICB treatment and the fitness function is defined as the accuracy of predicting a user defined phenotype (used as pseudo-labels) based on the total number of interactions from those selected occurring in a given sample; accuracy is quantified by the AUC. To minimize the risk of overfitting to the TCGA dataset and aid in faster convergence of the genetic algorithm during training, the size of the feature space is reduced by filtering out complex and harder to measure interactions involving receptors/ligands encoded by multiple genes and additionally assessing the fold change of each interaction between the two classes specified in the training dataset and incorporated into the fitness function (e.g., hypermutated vs non-hypermutated). After this step, only interactions with a fold change >1 is passed to the genetic algorithm for further optimization.

The algorithm starts out by randomly generating sets of features. This is defined as the seed population. These sets iteratively evolve via the phenomenon of natural selection enforced by the user defined fitness function. Specifically, for each subsequent iteration, features from the best performing sets, as determined by the user defined fitness function, in the current iteration are mixed at random followed by random new feature additions or dropouts (referred to as mutations) to build a new generation of feature sets and the process repeats. Eventually, after a number of epochs, which we set to 100, the fitness function converges to an optimum and the best set of features for the prediction task is returned to the user. Since the fitness function landscape is often non-convex and the training process is stochastic, we repeat the training process 500 times, each with a randomly chosen seed population, and eventually choose frequently selected features over all solutions to reach a solution we suspect is close to the global optimum solution and less likely to overfit. For our plots, we set the threshold to frequency > 100 times. The probability of any feature being selected more than 100 times by random chance based on this approach is estimated to be < 0.01. Results are qualitatively similar for more stringent thresholds. The genetic algorithm was implemented in R using the genalg package^72^

## Data availability

All deconvolved expression data generated in this study will be made publicly available upon publication.

## Code availability

The tools and all codes for reproducing the figures of this study will be made publicly available upon publication.

## Acknowledgements

This research is supported in part by the Intramural Research Program of the National Institutes of Health, National Cancer Institute, Center for Cancer Research. This work utilized the computational resources of the NIH HPC Biowulf cluster. We would additionally like to acknowledge Dr. Noam Auslander, Dr. Chi-Ping Day and other members of the Cancer Data Science Laboratory for their helpful feedback on this work.

## Author contributions

K.W., S.P. A.A.S. and E.R. conceptualized and designed the project. A.A.S. and E.R. supervised the project. K.W. and S.P. implemented the software and performed the data analyses. K.W., S.P., A.A.S. and E.R. wrote the original draft with input from all authors. J.L., E.M.G., W.R., F.S., and D.R.C. contributed to data analyses, visualization, review and editing.

