## Supplementary Note for "Deconvolving clinically relevant cellular immune crosstalk from bulk gene expression using CODEFACS and LIRICS"

#### Supplementary Figures S1-S12

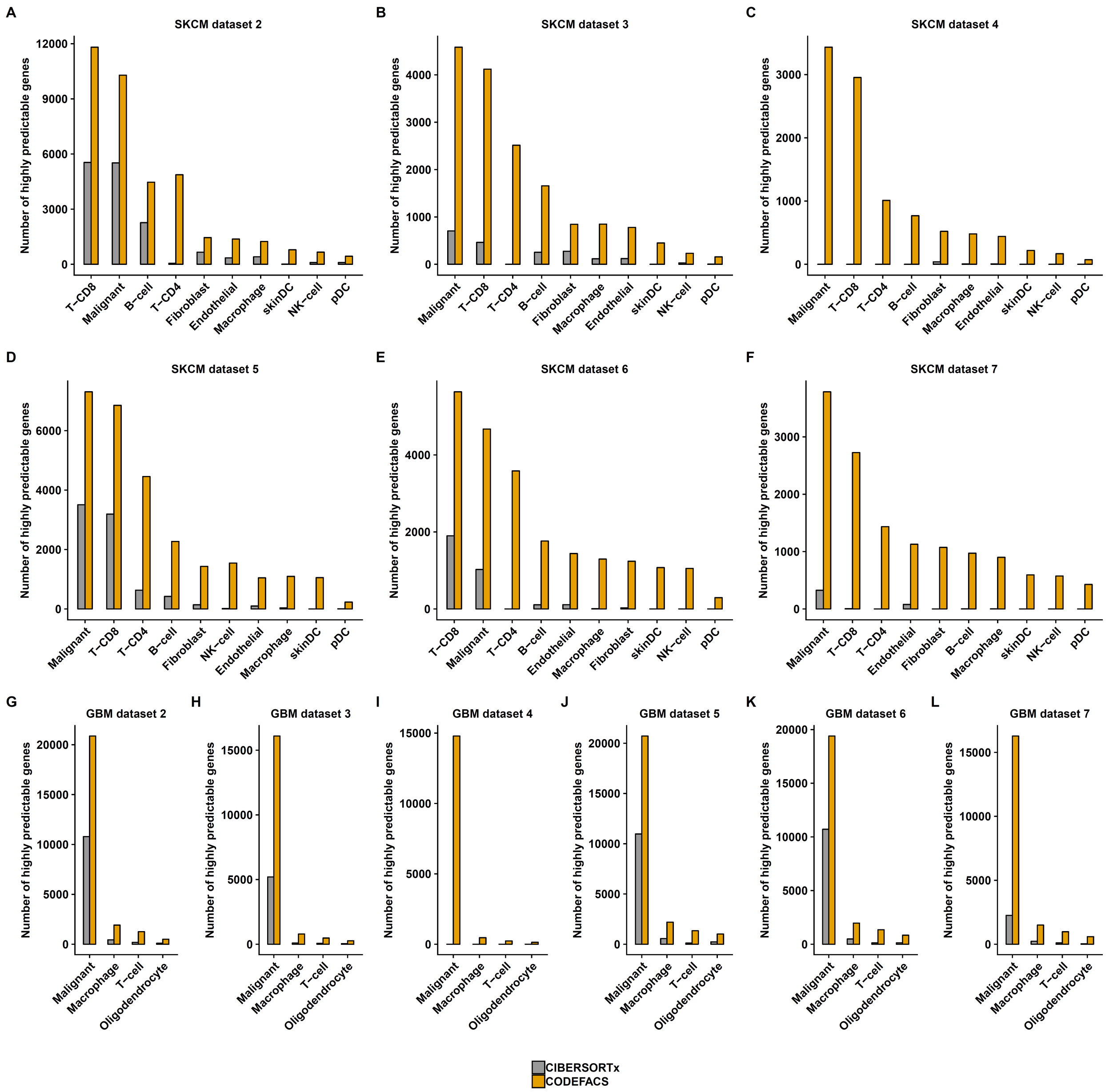

**Supplementary Figure S1: Number of highly predictable genes among 12 validation datasets for CIBERSORTx and CODEFACS. (A-L)** bar plots depicting the number of highly predictable genes (Kendall correlation ≥ 0.3) among 12 validation datasets (SKCM dataset 2, SKCM dataset 3, SKCM dataset 4, SKCM dataset 5, SKCM dataset 6, SKCM dataset 7, GBM dataset 2, GBM dataset 3, GBM dataset 4, GBM dataset 5, GBM dataset 6, GBM dataset 7). See **Supplementary Table 6** for the details of all the benchmark datasets. The yellow bar represents the performance of CODEFACS, while the gray bar represents that of CIBERSORTx.

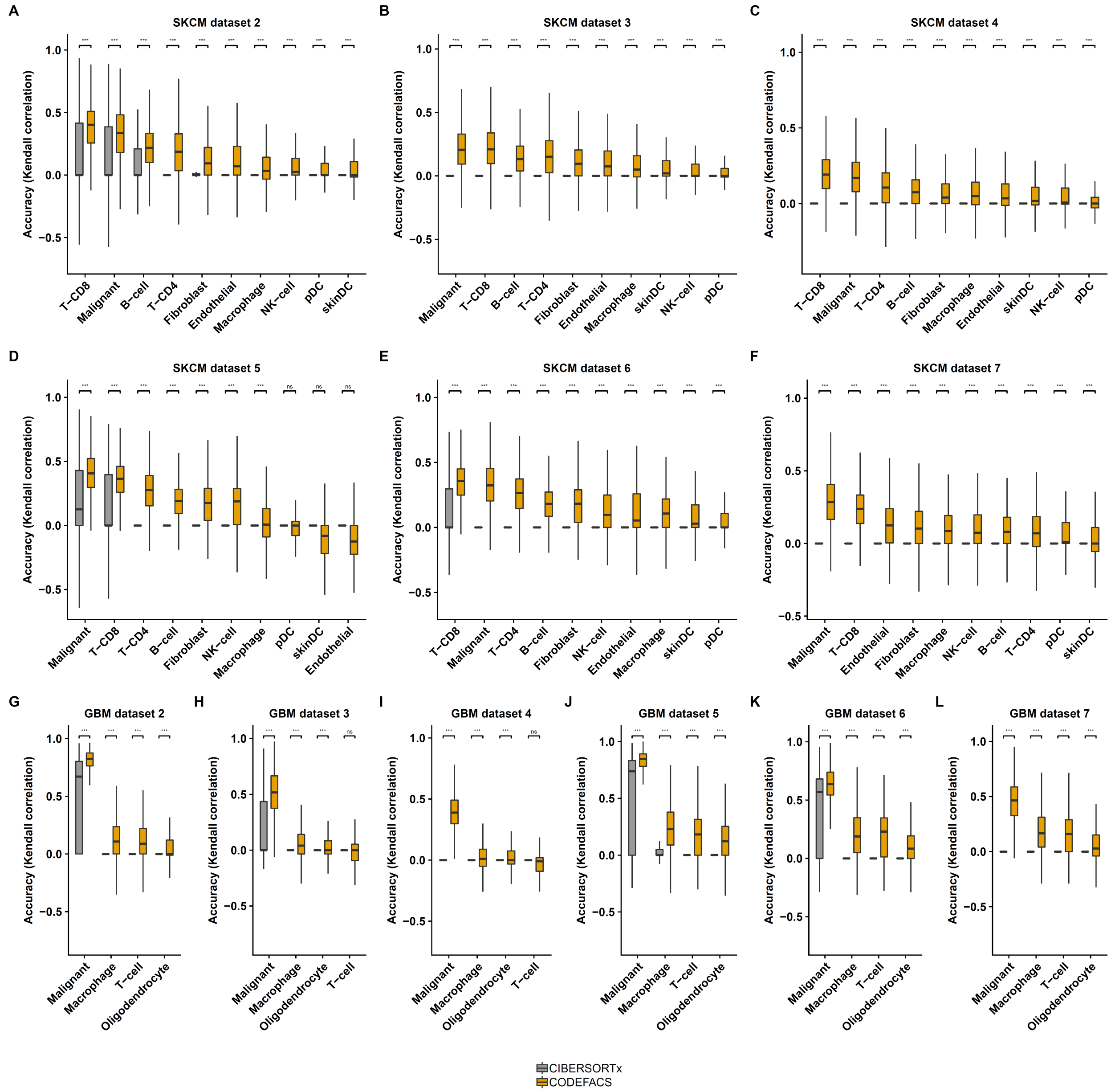

**Supplementary Figure S2: Accuracy (Kendall correlation) distribution comparisons among 12 validation datasets between CIBERSORTx and CODEFACS for high-confidence predictions (based on CODEFACS). (A-L)** boxplots depicting the accuracy distributions among the 12 benchmark dataset (SKCM dataset 2, SKCM dataset 3, SKCM dataset 4, SKCM dataset 5, SKCM dataset 6, SKCM dataset 7, GBM dataset 2, GBM dataset 3, GBM dataset 4, GBM dataset 5, GBM dataset 6, GBM dataset 7). See **Supplementary Table 6** for the details of all the benchmark datasets. The yellow box represents the performance of CODEFACS, while the gray box represents that of CIBERSORTx. Wilcoxon signed rank test was performed to compare the prediction accuracies of CODEAFCS and that of CIBERSORTx for each cell type in each dataset. *** denotes p-values < 2e-16.

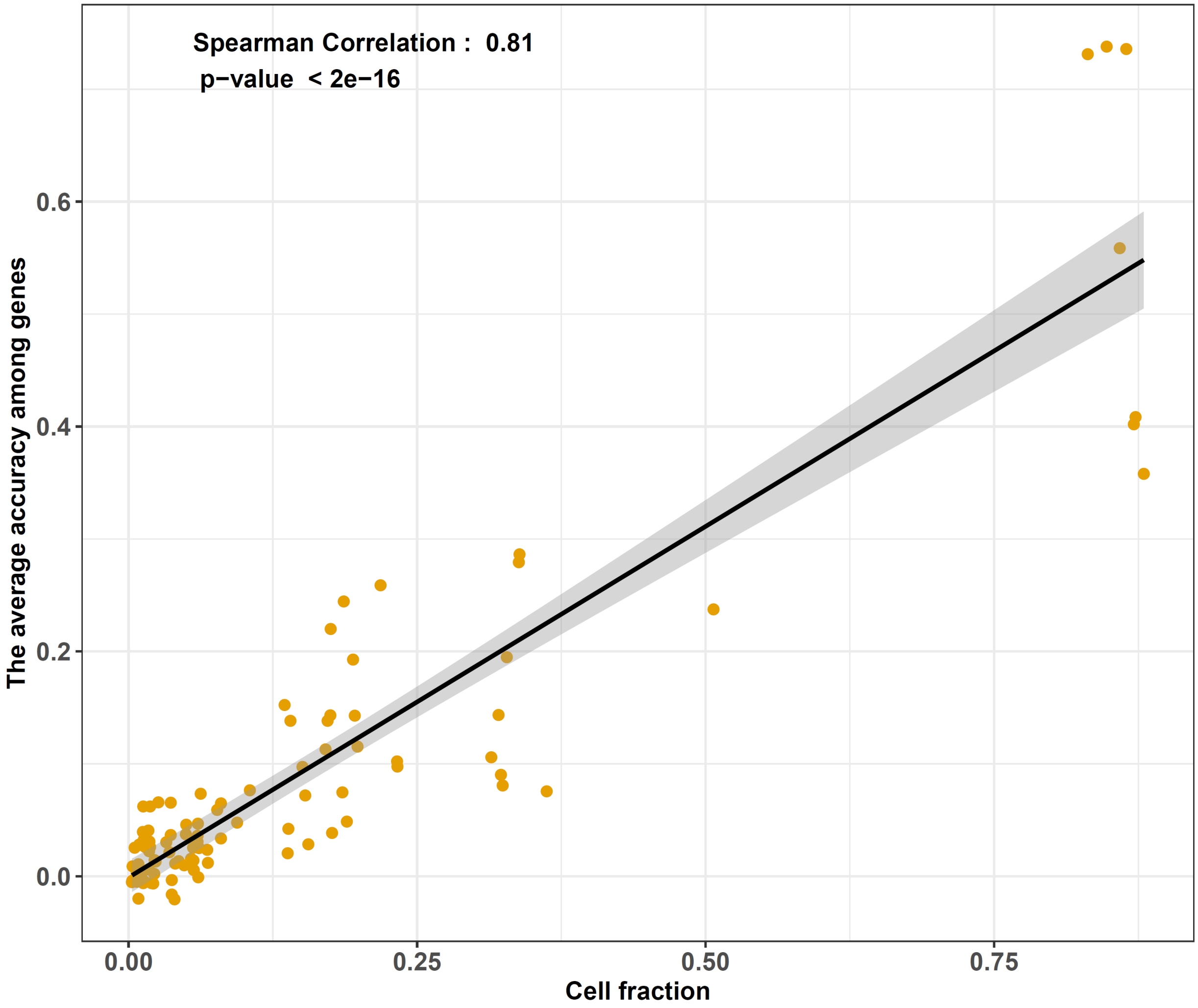

**Supplementary Figure S3: Scatter plot depicting the correlation between the average prediction accuracies (among genes) and cell type fractions across cell types and benchmark datasets for CODEFACS.** For each cell type of each dataset, the average prediction accuracy was computed by taking the average of prediction accuracies (Kendall correlation) across all genes. The y-axis indicates the average prediction accuracy among genes and the x-axis indicates the cell fraction. The Spearman correlation coefficient is ~0.81 (p-value < 2e-16).

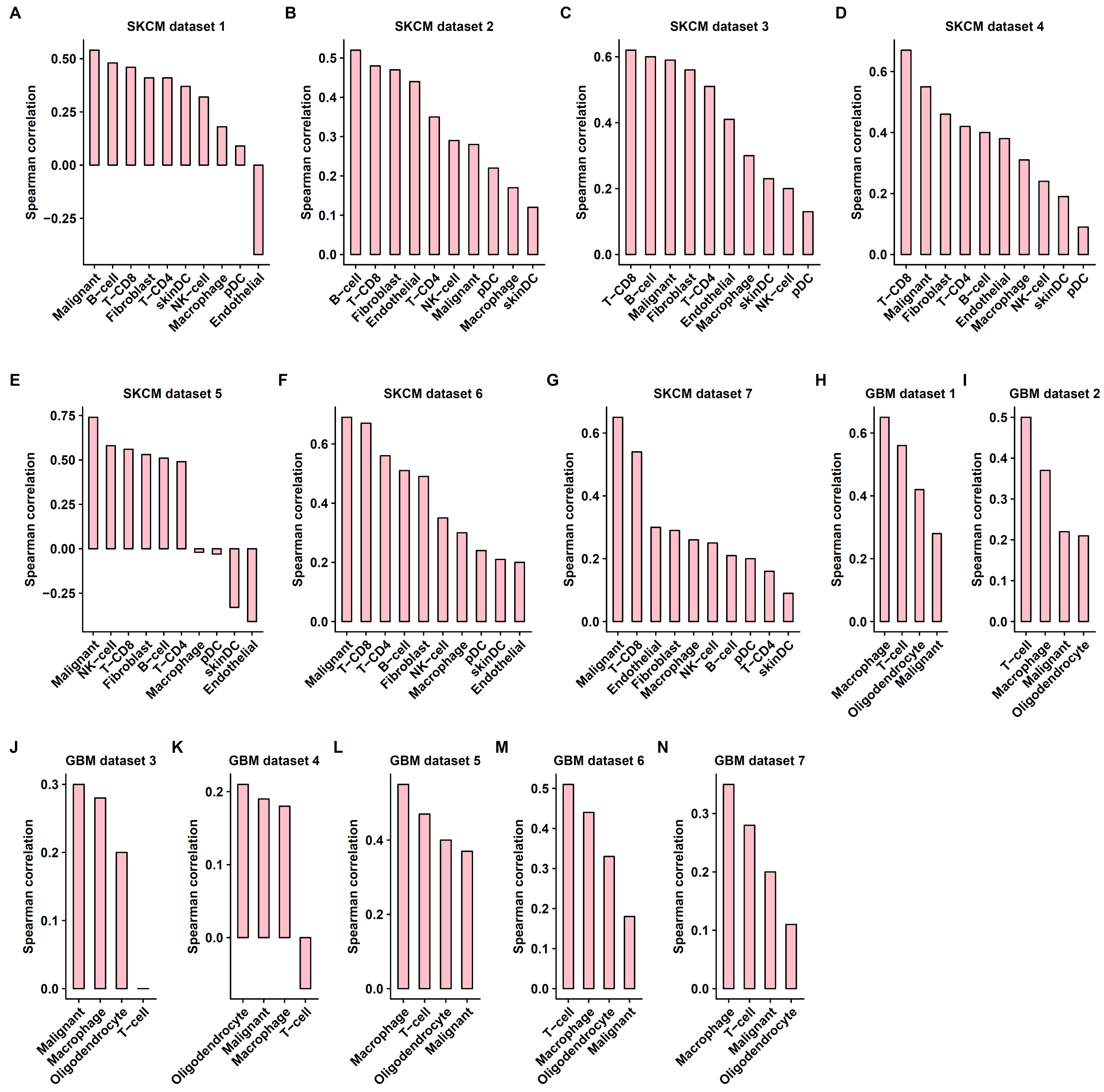

**Supplementary Figure S4: Correlations between prediction accuracies and confidence scores among cell types and benchmark datasets for CODEFACS. (A−N)** bar plots depicting the Spearman correlation between prediction accuracies and confidence scores in each cell type for all 14 benchmark datasets (SKCM dataset 1, SKCM dataset 2, SKCM dataset 3, SKCM dataset 4, SKCM dataset 5, SKCM dataset 6, SKCM dataset 7, GBM dataset 1, GBM dataset 2, GBM dataset 3, GBM dataset 4, GBM dataset 5, GBM dataset 6, GBM dataset 7). See **Supplementary Table 6** for the details of all the benchmark datasets. The y-axis indicates the Spearman correlation value, while the x-axis indicates the cell types.

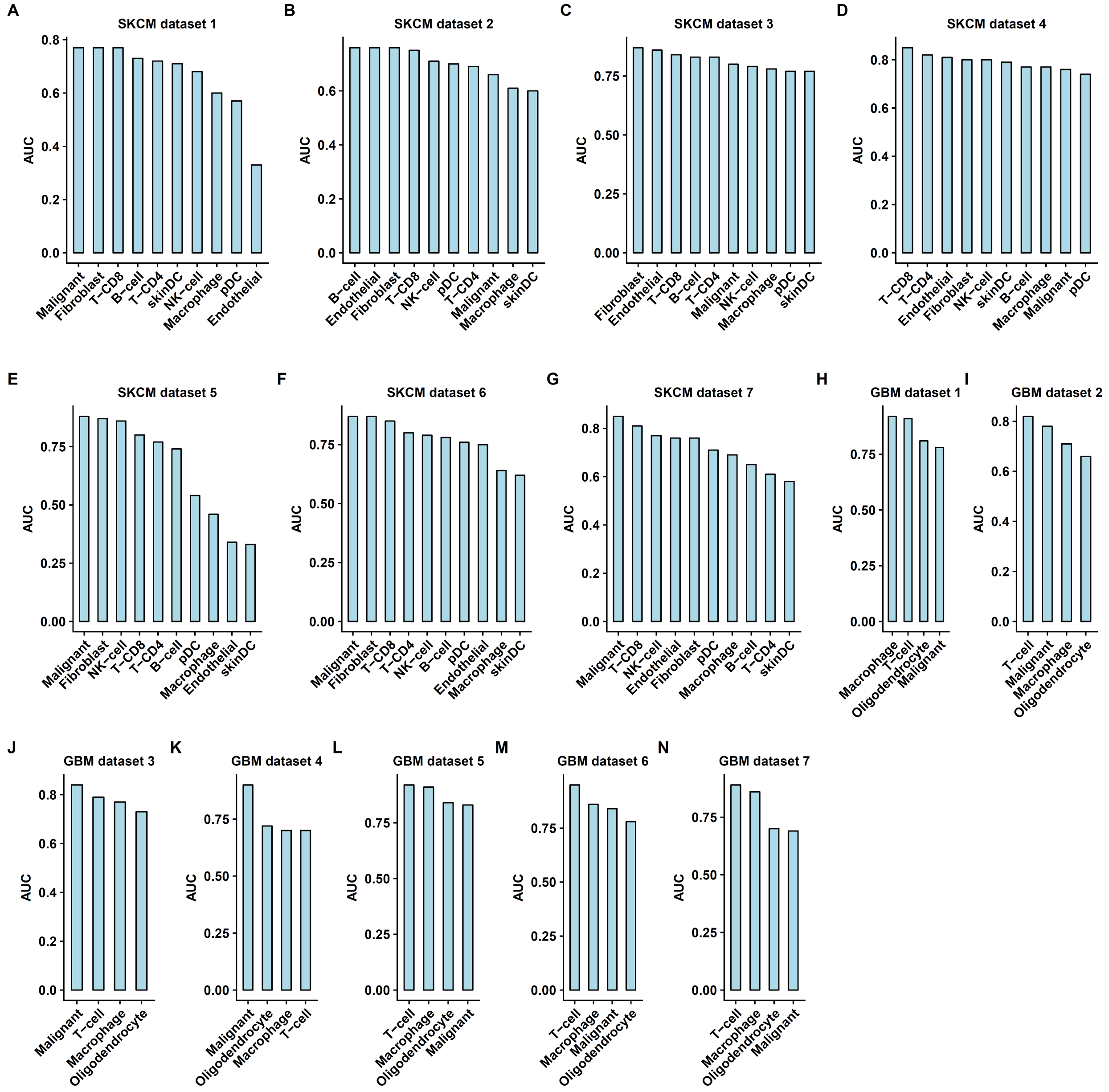

**Supplementary Figure S5: AUC for confidence scores in classifying informative and uninformative predictions among cell types and 13 benchmark datasets. (A−N)** bar plots depicting the AUC in each cell type for all 14 benchmark datasets (SKCM dataset 1, SKCM dataset 2, SKCM dataset 3, SKCM dataset 4, SKCM dataset 5, SKCM dataset 6, SKCM dataset 7, GBM dataset 1, GBM dataset 2, GBM dataset 3, GBM dataset 4, GBM dataset 5, GBM dataset 6, GBM dataset 7). Genes in each cell-type are grouped into two classes based on the correlation between their predicted and actual expression, informative (prediction accuracy ⩾ 0.1 and p-value ⩽ 0.05) and uninformative (prediction accuracy < 0.1 or p-value > 0.05). See **Supplementary Table 6** for the details of all the benchmark datasets. The y-axis indicates the AUC, while the x-axis indicates the cell type.

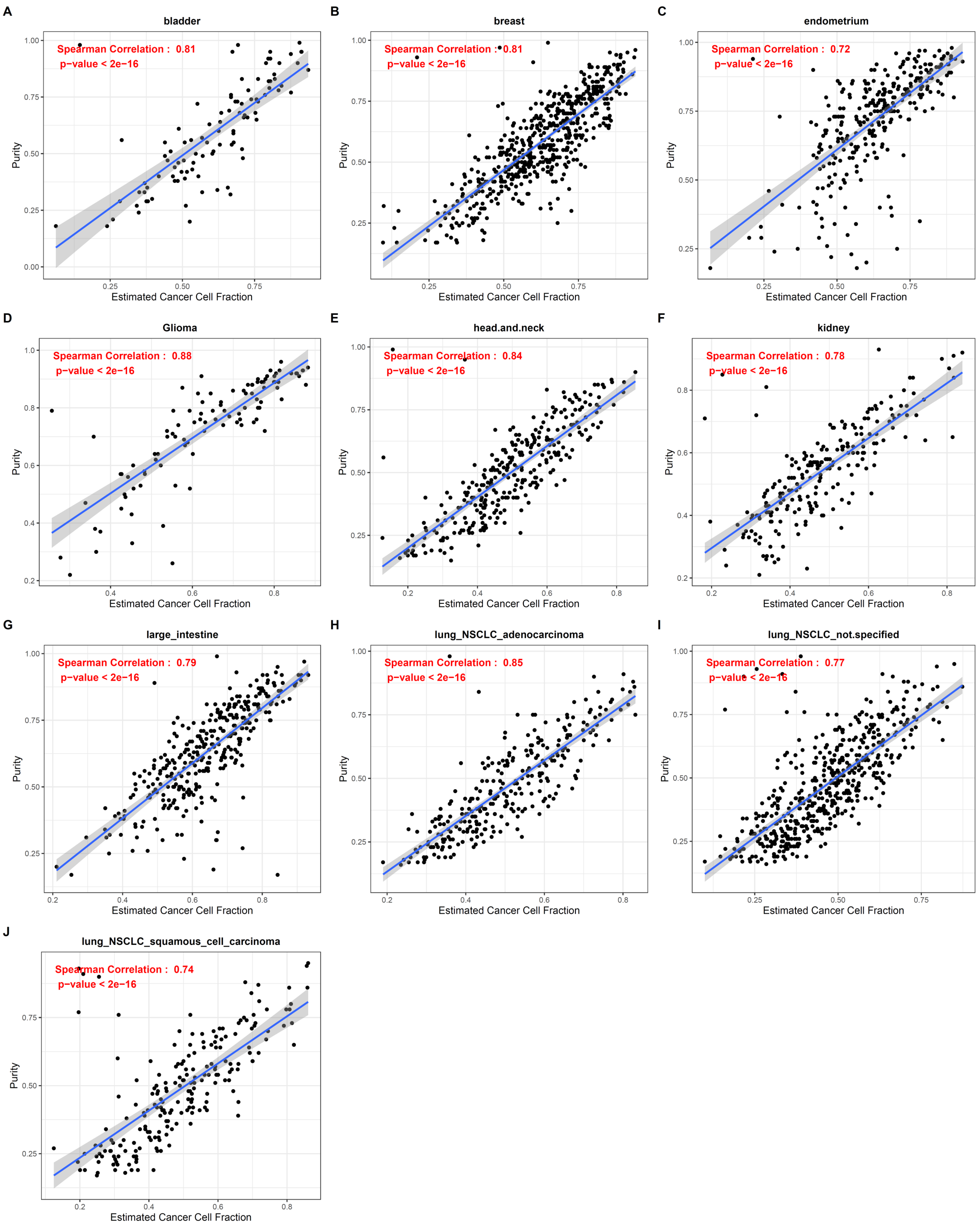

**Supplementary Figure S6: Correlations between predicted cancer cell fractions (by CODEFACS) and tumor purities (by ABSOLUTE) in TCGA. (A-J)** scatter plots depicting the correlation in bladder cancer, breast cancer, endometrium cancer, glioma, head and neck cancer, kidney cancer, large intestine cancer, lung adenocarcinoma, combined lung cancer, lung squamous cell carcinoma respectively. The y-axis denotes the reported purity in TCGA and the x-axis denotes the predicted cancer cell fraction. Both the Spearman correlation coefficient and the p-value are shown in the scatter plots.

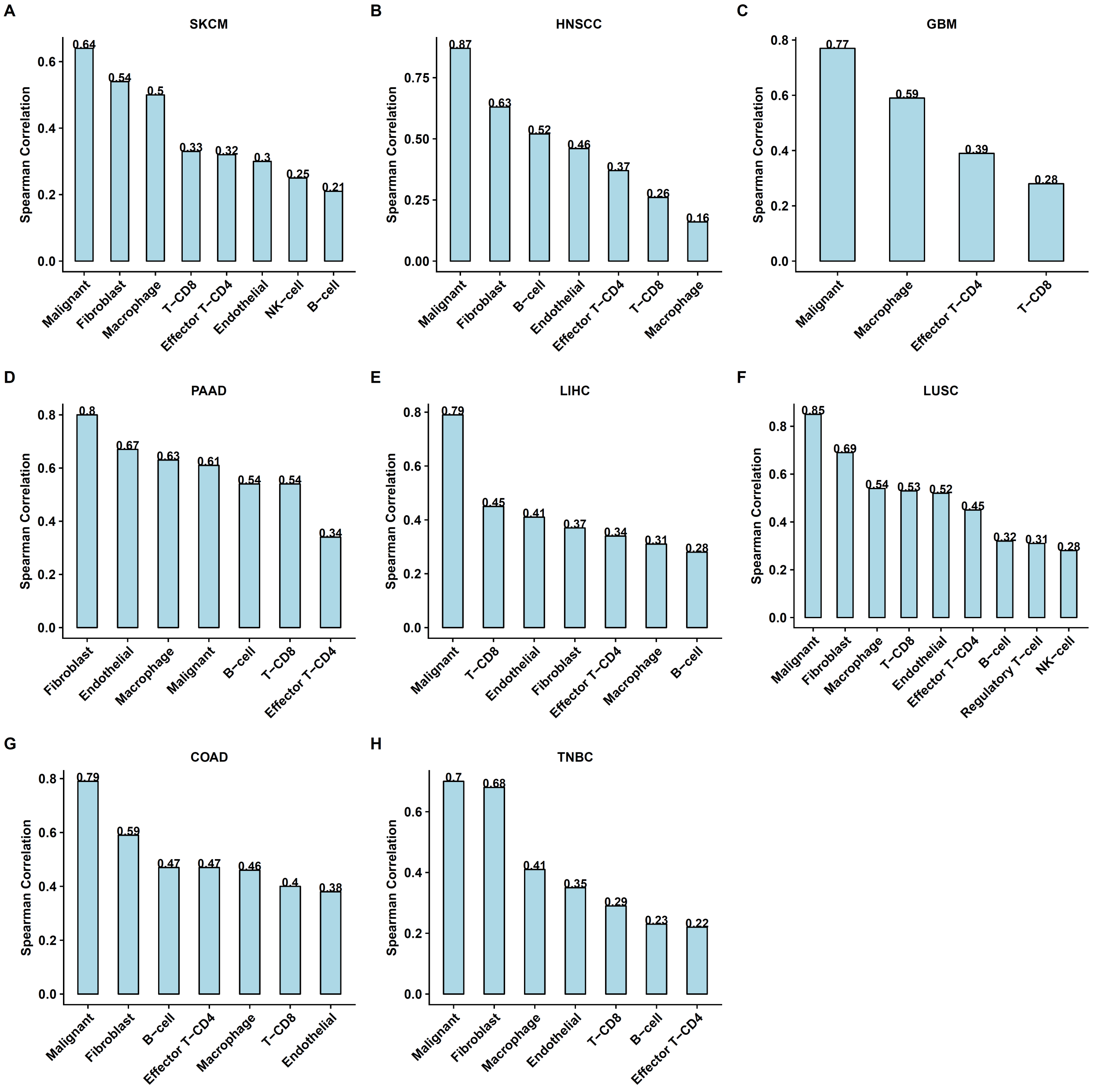

**Supplementary Figure S7: Correlation between estimated average gene expression in deconvolved TCGA and that in publicly available single cell datasets across genes among corresponding cell types and cancer types. (A-H)** bar plots depicting the result for SKCM, HNSCC, GBM, PAAD, LIHC, LUSC, COAD and TNBC respectively. The y-axis indicates the Spearman correlation coefficient value, and the x-axis indicates the cell type.

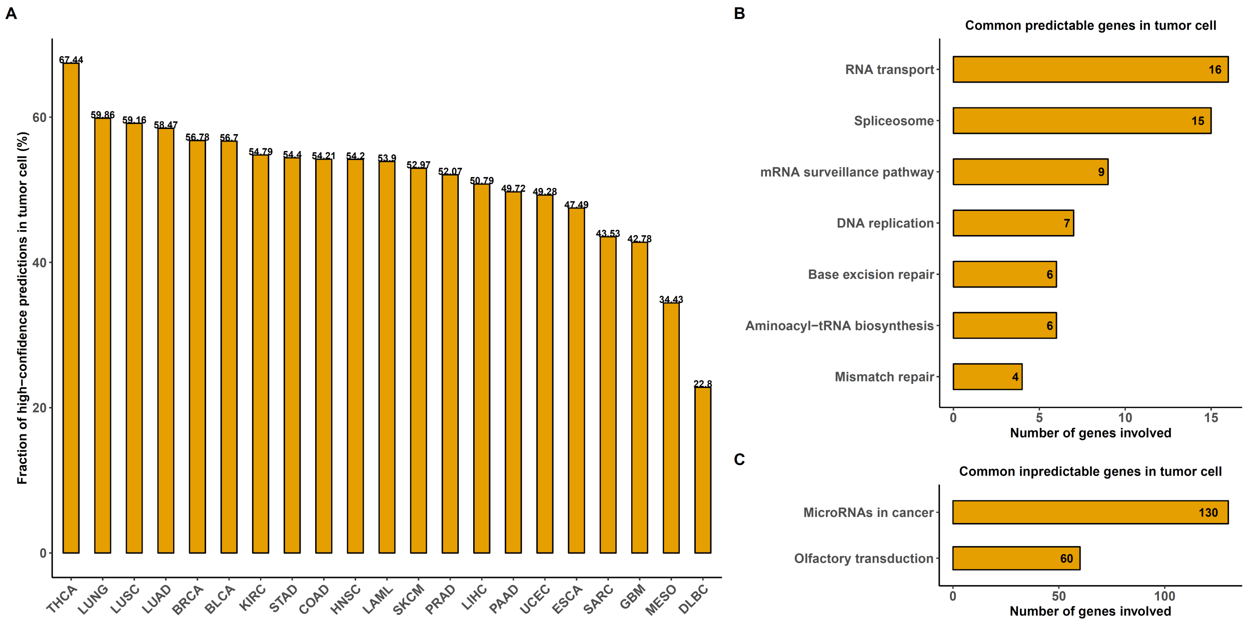

**Supplementary Figure S8: (A) bar plots depicting the fraction of predicted genes with high confidence in tumor cell among the 21 cancer types.** The y-axis indicates the fraction and x-axis indicates the cancer type; **(B) significantly enriched KEGG pathways among 325 genes commonly predicted with high confidence across 21 cancer types.** The y-axis indicates the enriched pathway and x-axis indicates the number of genes involved in each pathway among the 325 genes. **(C) significantly enriched KEGG pathways among 1733 genes commonly predicted with high confidence across 21 cancer types.** The y-axis indicates the enriched pathway and x-axis indicates the number of genes involved in each pathway among the 1733 genes.

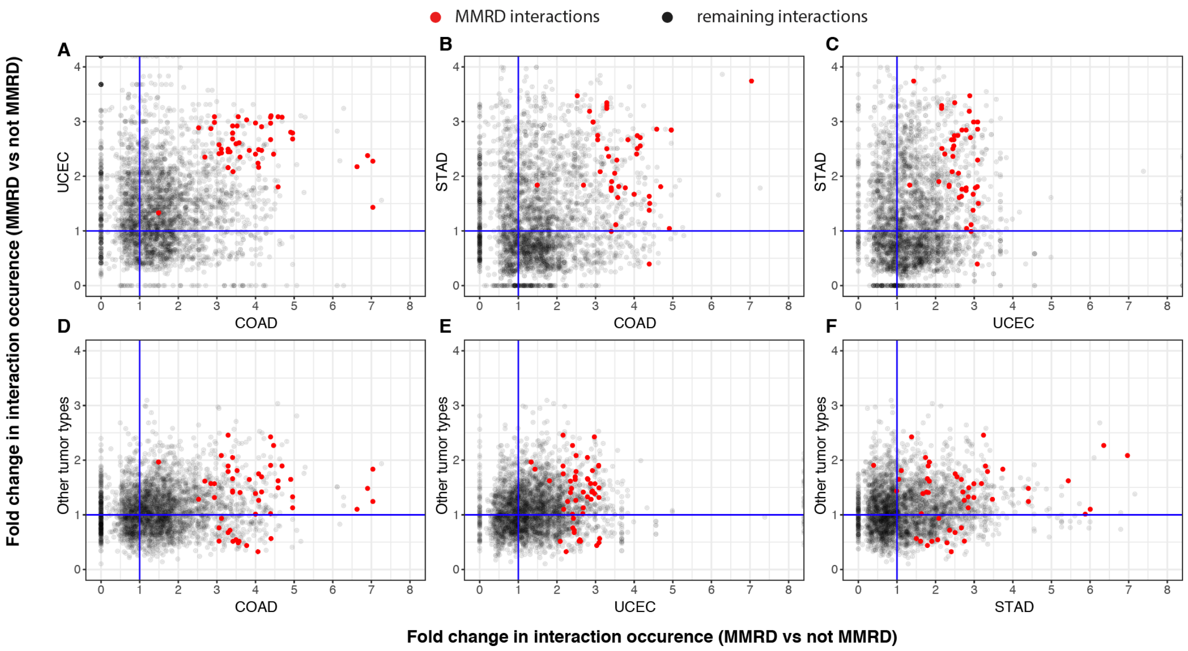

**Supplementary Figure S9: (A-F) a comparison of differential cellular crosstalk in mismatch repair deficient tumors from different tissues of origin.** Tumors are grouped into four different groups by tissue of origin: STAD (stomach), COAD (colon), UCEC (endometrium) and other (all other solid tumor types) in order to have sufficient numbers of mismatch repair deficient vs mismatch repair proficient samples per group. The axes measure the enrichment scores of all plausible ligand-receptor interactions between cell types in their respective group. A fold change > 1 implies the interaction occurs more frequently in mismatch repair deficient tumors. Interactions highlighted in red represent the shared core set of interactions from Figure 4A that are universally enriched in mismatch repair deficient solid tumors.

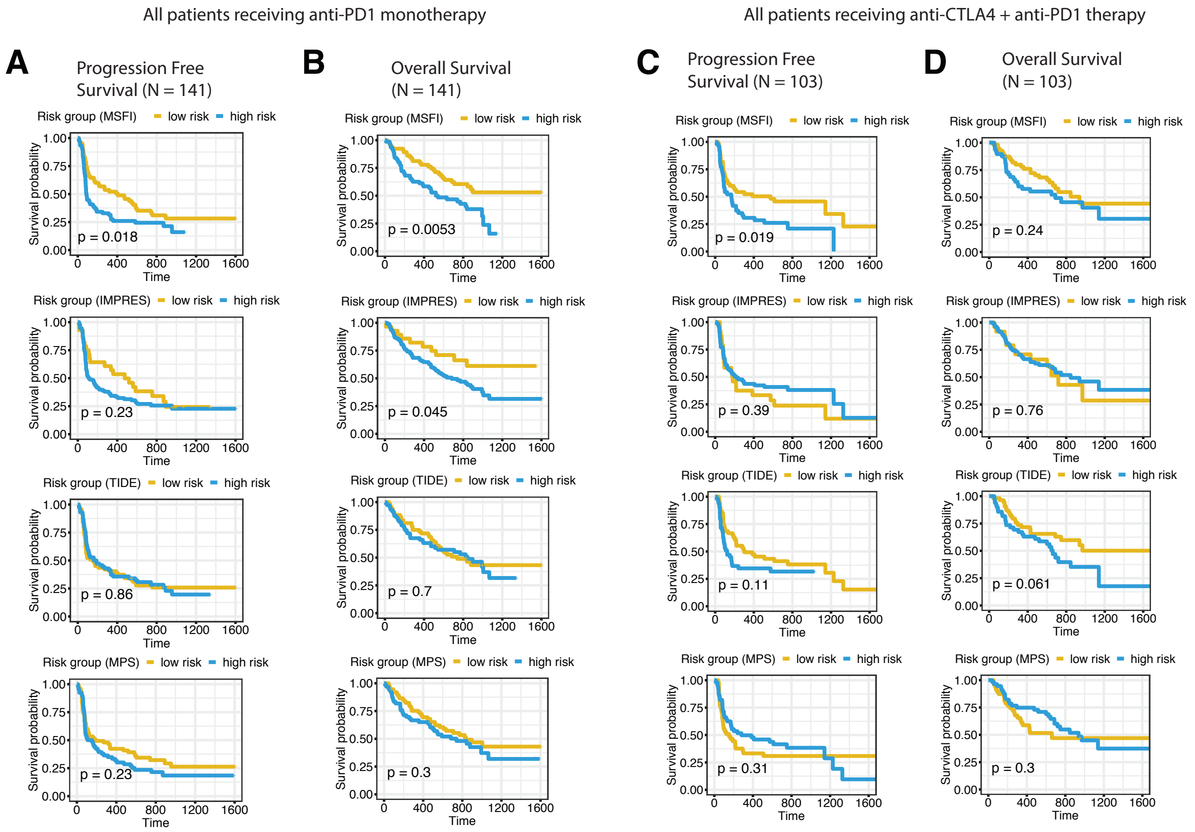

**Supplementary Figure S10: Comparison of Progression free survival and overall survival differences of patients receiving anti-PD1 monotherapy treatment vs anti-CTLA4 + anti-PD1 combination.** All patients receiving anti-PD1 treatment are classified into low-risk group if their LIRICS based cellular crosstalk score (MSFI score) exceeds the population median. The Likewise, when using the IMPRES score. For TIDE and MPS scores, all patients receiving anti-PD1 treatment are classified into low-risk group if their values fall below the population median (as these scores were shown to be associated with immune resistance[1,2]) **(A,B)** depict the survival differences for patients receiving anti-PD1 monotherapy only. **(C,D)** depict the survival differences for patients receiving anti-CTLA4 + anti-PD1 combination. Significance in the difference of survival trends of the two groups is calculated using the log-rank test. Time on the x-axis is measured in days.

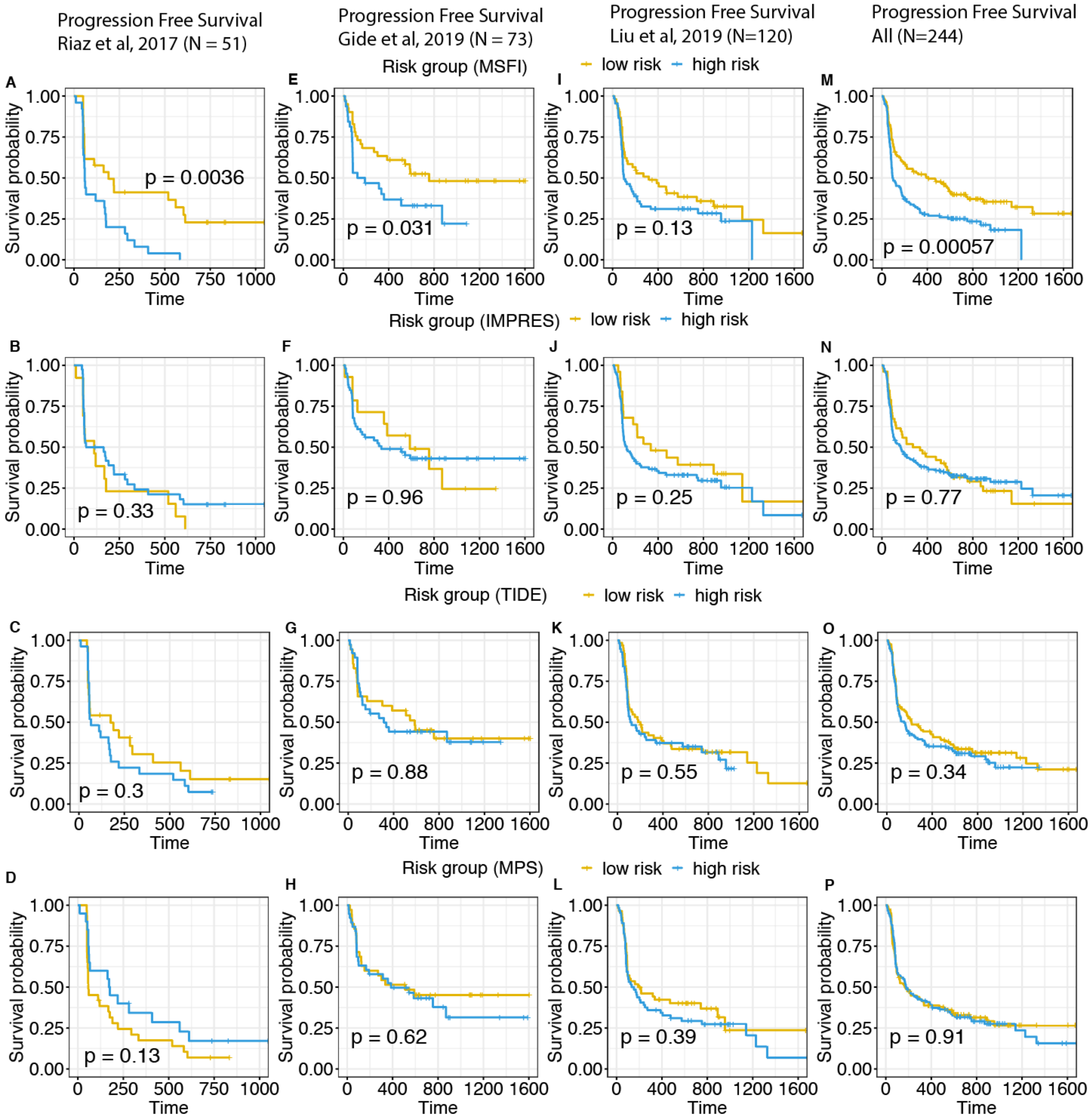

**Supplementary Figure S11: Progression free survival of patients receiving anti-PD1 treatment in each dataset [1–3]. (A, E, I, M)** Patients are classified into low-risk group if their LIRICS based cellular crosstalk score exceeds the population median. **(B, F, J, N)** Patients are classified into low-risk group if their IMPRES score [3] exceeds the population median. **(C, G, K, O)** Patients are classified into low-risk group if their TIDE score [1] is less than the population median. **(D, H, L, P)** Patients are classified into low-risk group if their MPS score [2] is less than the population median. Significance in the difference of survival trends of the two groups is calculated using the log-rank test. Time on the x-axis is measured in days.

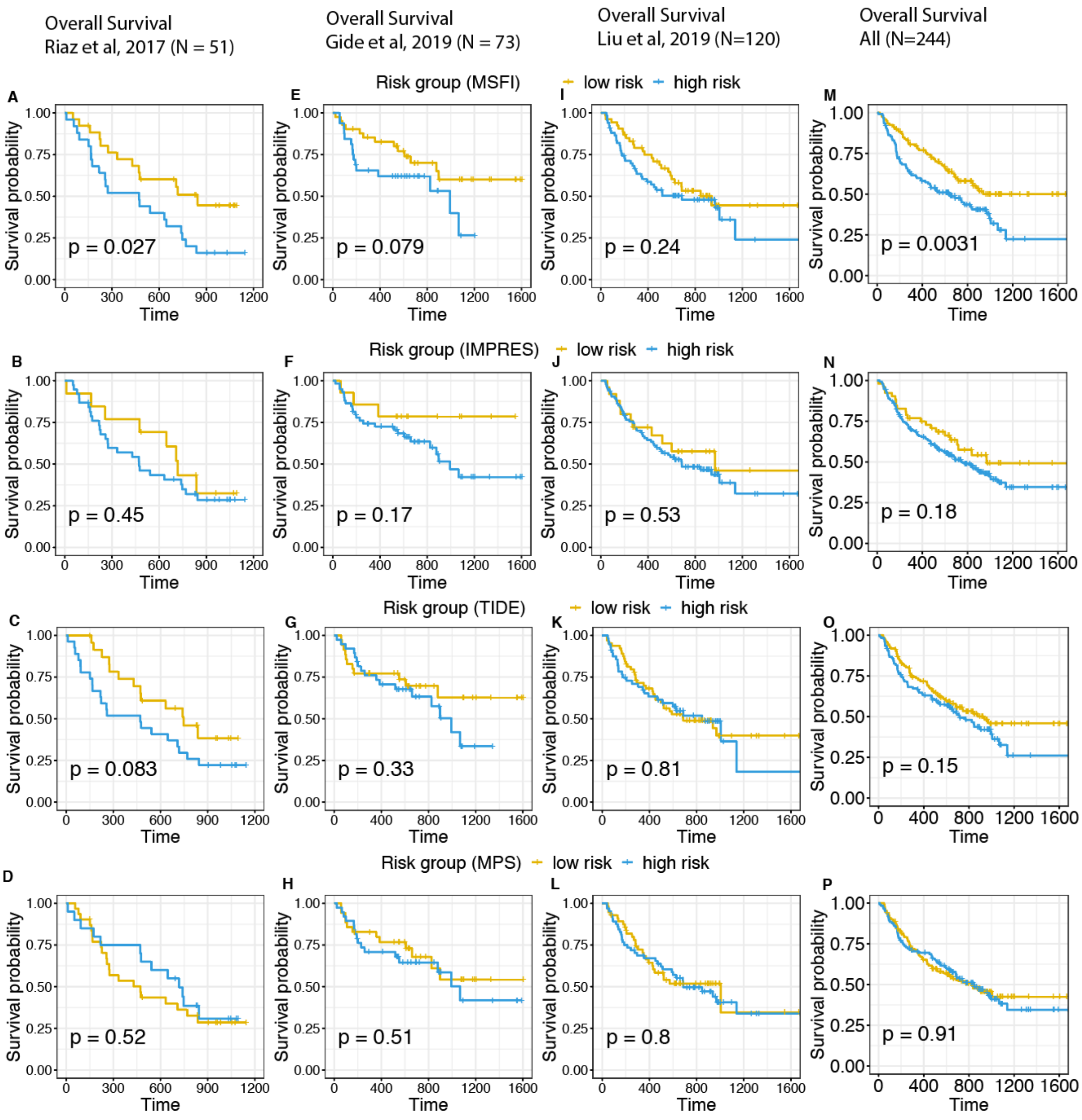

**Supplementary Figure S12: Overall survival of patients receiving anti-PD1 treatment in each dataset[1–3]. (A, E, I, M)** Patients are classified into low-risk group if their LIRICS based cellular crosstalk score exceeds the population median. **(B, F, J, N)** Patients are classified into low-risk group if their IMPRES score [3] exceeds the population median. **(C, G, K, O)** Patients are classified into low-risk group if their TIDE score [1] is less than the population median. **(D, H, L, P)** Patients are classified into low-risk group if their MPS score [2] is less than the population median. Significance in the difference of survival trends of the two groups is calculated using the log-rank test. Time on x-axis is measured in days.

#### Supplementary Note 1: CODEFACS (Confident DEconvolution For All Cell Subsets)

CODEFACS implements a divide-and-conquer strategy to reconstruct cell-type-specific transcriptomes from individual bulk mixtures *in silico*.

##### Input

1. Bulk gene expression of a collection of samples (required);
2. Cell fraction estimates of expected tumor, immune and stromal cell types in each sample (required if cell type specific signature profile is not provided); OR
3. Cell-type-specific signature profile (required if cell fractions are not provided).

##### Output

1. CODEFACS predicts the expression of each gene in each sample in each cell type in the mixture;
2. Confidence scores [0-1]for each gene-cell-type pair, which denote the confidence level in the predicted expression of a gene in a cell type across samples. (~1 High confidence, ~0 low confidence).

##### The full deconvolution problem

In this section, we provide a detailed description of the computational problem being solved by CODEFACS. The *full deconvolution* problem is formulated as follows[4]:

$$\text{minimize} \quad \sum_{i=1}^{m} ||(B_{i,\cdot} - \text{diag}(G_{i,\cdot,\cdot} \times F^T)||$$
$$\text{subject to} \quad \sum_{k=1}^{c} f_{jk} = 1 \text{ for all } j$$
$$g_{ijk} \geq 0 \text{ for all } i, j, k$$
$$f_{jk} \geq 0 \text{ for all } j, k \qquad (1)$$

where $B$ represents the given bulk RNA-seq expression matrix ($m$ genes x $n$ samples), in which each entry $b_{ij}$ is the observed bulk expression for $i^{\text{th}}$ gene and $j^{\text{th}}$ sample; $G$ is a three-dimensional deconvolved expression matrix ($m$ genes x $n$ samples x $c$ cell types), in which $g_{ijk}$ denotes the unknown expression for gene $i$ in the $j^{\text{th}}$ sample and $k^{\text{th}}$ cell type; $F$ is the cell fraction matrix ($n$ samples x $c$ cell types), in which $f_{jk}$ denotes the unknown cell fraction of $k^{\text{th}}$ cell type in $j^{\text{th}}$ sample. $F$ varies across samples and cell types but is constant across genes. $||\cdot||$, represents the $L_2$-norm (which measures the reconstruction error) and $\text{diag}()$ represents a function that gives a vector by extracting the diagonal entries of a matrix. The objective is to find an optimal solution for $G$ and $F$ with the constraint that the cell fractions (of $c$ cell types) in any sample $j$ sum up to 1 and all the gene expression values $g_{ijk}$ are non-negative real values. In this study, we assume the gene expression is quantified as TMM normalized TPM values. More specially, we employed a strategy introduced by Monaco et al. [5], which first estimates between-sample scaling factors upon raw TPM values using TMM method [6] and further scale TPM values in each sample using these scaling factors.

Problem (1) has no unique optimal solution without additional constraints and regularizations since there are more parameters to be estimated than observations[4,7]. However, problem (1) can be separated into two independent problems: cell fraction estimation and cell-type-specific gene expression prediction for each individual sample. The cell fraction estimation problem is formulated as follows:

$$\text{minimize } \sum_{i=1}^{l} ||B'_{i,\cdot} - S_{i,\cdot} \times F^T||$$
$$\text{subject to } \sum_{k=1}^{c} f_{jk} = 1 \text{ for all } j,$$
$$f_{jk} \geq 0 \text{ for all } j, k \qquad (2)$$

Where $S$ denotes the cell-type-specific signature matrix ($\ell$ genes x $c$ cell types) and the $\ell$ genes are a subset of all the $m$ genes in $G$ or $B$ matrix that are preferentially over-expressed in at least one of the $c$ cell types and their expression is assumed to be constant across the population to arrive at an approximate solution of cell fractions in each sample. $F$ is the same as that in equation (1), while $B'$ is a submatrix of the bulk expression matrix $B$ in equation (1) corresponding to the $\ell$ genes in cell-type-specific signature matrix $S$. Numerous effective cell fraction estimation tools have been developed and reported to solve problem (2) [8–24]. The experimental analog to these methods is the cell gating procedure described in FACS. We solve this problem using a well-known reference-based approach--CIBERSORT [21]. If needed, CODEFACS provides a batch correction approach introduced by CIBERSORTx that could be applied to minimize cross-platform technical batch effects between bulk mixture profile and cell type signature profile generated from different technical platforms (e.g. bulk RNA sequencing, SmartSeq2-based single cell sequencing, 10x-based single cell sequencing and microarray expression profiling) [4]. In addition, we provide the option to input prior known cell fractions instead of performing cell-fraction estimation de novo. Either known cell fractions or cell type signature profiles are required as input. Newman et al. [4,21] found that cell fractions determined by the CIBERSORT algorithm, which we reimplement in CODEFACS, mostly exhibit strong concordance with ground truth.

Once $F$ is estimated or provided, the full deconvolution problem formulated in (1) can be reduced to solving for $G$, given $B$ and $F$. One can additionally reduce problem (1) to a simpler problem where one solves for the expected cell-type-specific expression for a specific gene across a group of individual samples, given the cell fractions and bulk expression matrix:

$$\text{minimize } \sum_{i=1}^{m} ||B_{i,\cdot} - \bar{E}_{i,\cdot} \times F^T||$$
$$\text{subject to } \bar{e}_{ik} \geq 0 \text{ for all } i,k \; (3)$$

where $B$ is the same as in equation (1) and represents the input bulk expression matrix; $F$ is also the same as that in equation (1) and denotes cell fractions; $\bar{E}$ is the expected cell-type-specific expression matrix ($m$ genes x $c$ cell types) across the population, in which $\bar{e}_{ik}$ denotes the expected expression of gene $i$ in cell type $k$. For a fixed $F$, a unique optimal solution for this problem exists and can be found using non-negative least squares (NNLS) [4,25,26]. The key difference between problem (3) and problem (1) is that the former aims to predict expected cell-type-specific expression for each gene in the population, while the latter predicts cell-type-specific expression for each gene and each sample.

One can aim to solve problem (1) approximately by making use of a greedy divide and conquer strategy that breaks down problem (1) into simpler problems (2) and (3). Newman et al, in their groundbreaking work CIBERSORTx, were the first to propose such an algorithm. In CODEFACS, we introduce the concept of confidence scores and additional algorithmic improvements. We show that CODEFACS yields a much more accurate solution compared to CIBERSORTx in 15 benchmark datasets with ground truth data.

The CODEFACS algorithm consists of three modules that are executed sequentially and a confidence ranking system that is invoked after the execution of each module. In module 1 we refined and extended the high-resolution deconvolution module introduced by CIBERSORTx. First, we generalized their two-freedom estimation method into a recursive splitting method, which we call "$p$-freedom estimation" (The degrees of freedom represent the distinct latent sources of variability in gene expression across individuals). We found that $p$-freedom estimation could capture tumor heterogeneity better than the 2-freedom estimation. Second, we generalized their sliding window method by employing an ensemble of window sizes. Using an ensemble of window sizes seeks to reduce the dependence of downstream biological analyses on arbitrary choices of the window size parameter. In addition, we developed modules 2 and 3 (hierarchical deconvolution and imputation-based deconvolution) to further increase the number of highly predictable genes. The confidence ranking system uses a series of heuristics to decide where the solution can be improved by subsequent modules. See Figure 1A for the schematic diagram with the inputs and outputs.

**The notion of confidence**

Before we formally describe the algorithm, we introduce the concept of confidence, which is a central part of the algorithm. Each of the three prediction modules operates under specific modeling assumptions that are, in theory, uniformly applicable to all genes. However, in practice, certain genes might violate these assumptions. Therefore, for such genes, one cannot confidently say whether their predicted cell-type specific expression levels closely reflect the ground truth. To quantify this uncertainty, we designed a *confidence ranking system,* which can decide whether a specific prediction requires further refinement in subsequent modules by defining a ranking $\Phi$ over genes for each cell type using confidence relevant features (more details are provided subsections 3, 5, 7). Additionally, the confidence ranking system also re-evaluates the confidence level of each final prediction (gene-cell type pair) and provides in the end report a confidence score between 0 and 1 (section 8).

#### The CODEFACS algorithm introduced according to its workflow in the following sections

##### 1. Cell fractions estimation (optional)

###### 1.1 Cell fractions are estimated using SVM regression given cell type signatures and bulk expression (optional)

Cell type signatures are derived based on prior reference datasets using the signature derivation module from CIBERSORTx. Thereafter, we implemented a support vector machine (SVM)-regression-based method to predict cell fraction given bulk expression/methylation and prior cell-type-signature profiles following the CIBERSORT algorithm[21]. Given the bulk mixture and cell type signature profile, the SVM regression model outputs predicted cell fraction for each cell type and sample (**Supplementary Fig. S12**). If the user provides prior known cell fractions as input, CODEFACS will skip this optional step.

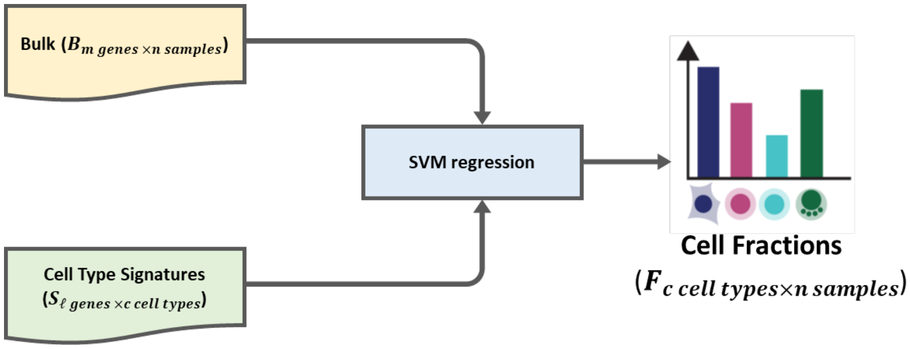

**Supplementary Figure S12: Cell fraction prediction using SVM regression.** Given the bulk mixture matrix $B\ (m\ \text{genes} \times n\ \text{samples})$ and cell type signature profile $S\ (\ell\ \text{genes} \times c\ \text{cell types})$, we use SVM regression to predict the cell fraction matrix $F\ (c\ \text{cell types} \times n\ \text{samples})$.

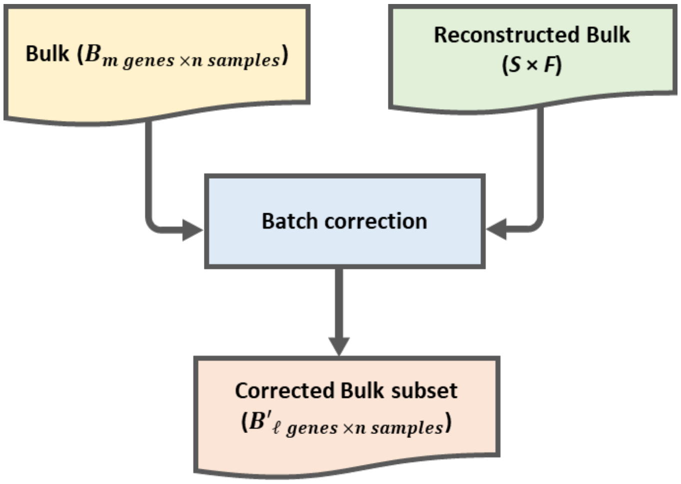

**Supplementary Figure S13: Batch correction module.** Given initial estimates of cell fractions $F\ (c\ \text{cell types} \times n\ \text{samples})$ and cell type signature profile $S\ (m\ \text{genes} \times \text{c cell types})$, one can reconstruct the bulk expression matrix via the matrix multiplication $S \times F$. Batch effects are then reduced between the given bulk subset $B'$ and reconstructed bulk $S \times F$.

##### 1.2 Batch correction to refine cell fractions (optional)

To account for any systematic batch effects between bulk expression and independently generated cell-type-specific signature expression data, which could bias cell-fraction estimates, we re-implemented the batch correction method introduced by CIBERSORTx [4]. The rationale for this method is that any batch effect between the given bulk expression and independently generated cell-type-specific signature expression must also be reflected in the reconstructed bulk expression $S \times F$. Thus, one can further refine cell-fraction estimates of each cell type in each sample after reducing the batch effect between the given bulk matrix and reconstructed bulk matrix $S \times F$ , using the function ComBat() from the SVA package [27] in R (**Supplementary Fig. S13**). The final output of this step is a refined cell-fraction matrix. Currently our implementation focuses on correcting biases among bulk RNA sequencing and SmartSeq2-based single cell sequencing datasets. This step is optional and will be skipped if the user does not specify that it should be done. For more details on the batch correction procedure, please refer to section “Cross-platform normalization schemes for deconvolution” in the supplementary information of CIBERSORTx[4].

#### 2. Module 1 - High resolution deconvolution

In this module, the observed bulk expression of a gene in a sample is modeled as the weighted sum of cell-type-specific expression of that gene from that sample (See problem 1 above).

##### 2.1 Determine cell types in which a specific gene is weakly expressed

To determine if a gene $i$ is weakly expressed in a cell type, we first conduct the following statistical analysis: individuals are randomly chosen without replacement to generate 100 random subsets of individual samples and then problem (3) is solved to estimate expected cell-type-specific expression for each random subset. This bootstrapping procedure generates a distribution of expected cell-type-specific expression values $\bar{e}_{ik}$ in the population. We then derive two p-values for each cell type $k$: first, an empirical p-value that is estimated by checking the percentage of solutions where $\bar{e}_{ik}$ > 0, and second, a p-value derived from a parametric t-test. The two p-values are then combined using Fisher’s method[28] to obtain a final p-value for each cell type. If a gene is weakly expressed in a cell type (FDR > 0.2), we force the cell fractions of that cell type in the corresponding mixture model to be 0 to improve the deconvolution of gene expression in other cell types.

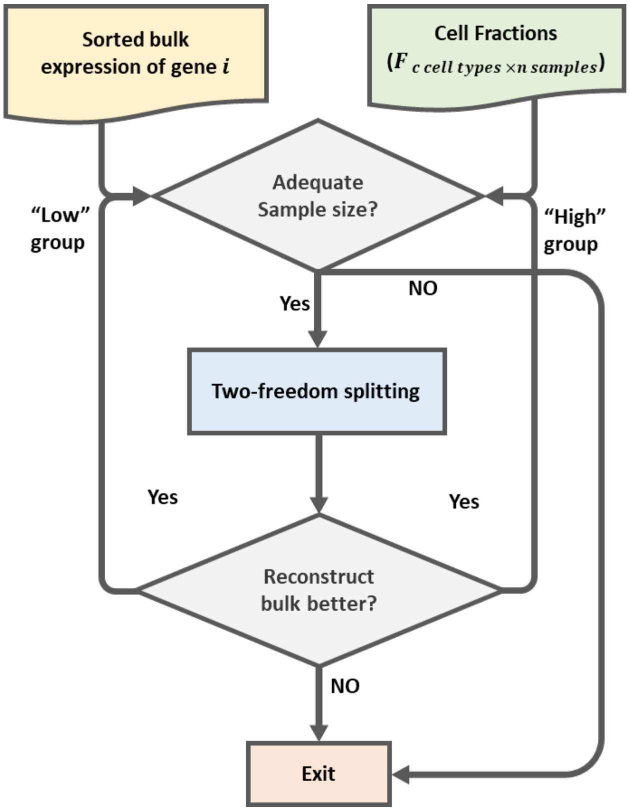

**Supplementary Figure S14: Recursive splitting method.** Given the estimated cell fractions $F$ $(c \text{ cell types } \times \ n \text{ samples})$ and sorted bulk expression of gene $i$, we check whether we have an adequate sample size for NNLS first: if not, it will exit; if yes, two-freedom splitting will be performed. Subsequently, we will check whether the two-freedom splitting improves the bulk reconstruction. If yes, both the low-expressed and high-expressed groups will recursively enter another round of two-freedom splitting; if no, the two-freedom splitting based predictions will be ignored and the function exits.

#### 2.2 Recursively splitting samples into finite sub-groups

With an appropriate cell-type-mixture model defined for each gene, we now try to find an approximate solution to problem (1). For a gene $i$, one can divide problem (1) into a finite number of simpler problems by assuming that individuals with similar bulk expression levels of gene $i$ must have similar cell-type-specific expression levels of gene $i$. Hence, we first sort all samples in increasing order according to the bulk expression of gene $i$. In the two-freedom deconvolution in CIBERSORTx algorithm, one can then find a position $t$ to partition all the sorted samples into two sorted subsets: $h_1 = \{1, 2, \dots, t-1\}$ and $h_2 = \{t, t+1, \dots, n\}$, such that the expected cell-type-specific expression in each of the subsets (obtained from solving problem 3 using NNLS) best reconstructs the observed bulk expression (For more details, see section "Cell type expression coefficients that best explain the bulk GEP" in the supplementary information of CIBERSORTx [4]). Either of these two subsets can now be recursively partitioned further into smaller subsets in a similar fashion if the re-construction error keeps dropping and the subsets sample size stays above 1.9 times the number of cell types. This is referred to as $p$-freedom approach which extends the two-freedom approach of CIBERSORTx (**Supplementary Fig. S14)**. The recursive splitting pseudo-code is shown below:

**recursive_splitting():**

**Input:**

*$F$= cell fraction matrix*

*$B$ = bulk expression matrix*

*$size$ = the number of samples required for NNLS*

*$i$ = gene index*

*$st$ = start position in the sorted list of samples*

*$ed$ = end position in the sorted list of samples*

*$\bar{E}_{i,\cdot}$ = expected cell-type-specific expression of gene i over all sorted samples in the range $\{st, \dots, ed\}$ (Obtained from solving problem 3)*

**Output:**

*$G^p_{i,\cdot,\cdot}$ = cell-type-specific expression over samples (1, 2, …, n) from recursive splitting deconvolution*

***Function:***

*if ($ed - st$ + 1 < 2 × $size$) {*

*save $\bar{E}_{i,\cdot}$ to $G^p_{i,\cdot,\cdot}$;*

*exit;*

*}*

*else {*

$t$ *= 2-freedom index that splits sorted samples in the range* $\{st, \ldots, ed\}$*, into two subsets:* $\{st, \ldots, st+t\}$ *and* $\{st+t+1, \ldots, ed\}$ *(See CIBERSORTx algorithm for more details)*

$\overline{L}_{i},.$ *= expected cell-type-specific expression of gene i over all sorted samples in the range* $\{st, \ldots, st+t\}$ *(Obtained from solving problem 3)*

$\overline{H}_{i},.$ *= expected cell-type-specific expression of gene i over all sorted samples in the range* $\{st+t+1, \ldots, ed\}$ *(Obtained from solving problem 3)*

*Err1 =* $\left\|B_{i,\{st,\ldots,ed\}} - \left(\overline{E}_{i},. \times \left(F_{\{st,\ldots,ed\}},.\right)^{T}\right)\right\|$

*Err2 =* $\left\|B_{i,\{st,\ldots,ed\}} - \left[\left(\overline{L}_{i},. \times \left(F_{\{st,\ldots,st+t\}},.\right)^{T}\right), \left(\overline{H}_{i},. \times \left(F_{\{st+t+1,\ldots,ed\}},.\right)^{T}\right)\right]\right\|$

*if (Err2 < Err1) {*

*Save* $\overline{L}_{i},.$ and $\overline{H}_{i},.$ *to* $G_{i}^{p},.,.;$

*Call: recursive_splitting(F, B, size, i, st, st+t,* $\overline{L}_{i},.$*);*

*Call: recursive_splitting(F, B, size, i, st+t+1, ed,* $\overline{H}_{i},.$*);*

*Exit;*

*}*

*Exit;*

*}*

***End***

##### 2.3 Ensemble sliding window deconvolution

For gene $i$, a sliding window is defined over the sorted list of samples with a specific window size $s$. For each window of sorted samples, problem (3) is solved using NNLS to estimate the expected cell-type-specific expression across samples within that window. Cell-type-specific expression for each individual is then approximated by redistributing population-level estimates of cell-type-specific expression within each sliding window (This is again based on the assumption that subsets of individuals with very similar bulk expression profiles have a shared cell-type-specific expression profile. For more details on how this is done, please refer to the CIBERSORTx algorithm[4]). Thereafter, the initial approximate predictions from the sliding window deconvolution of window size $s$ are refined using a linear-regression-based smoothing procedure such that the distribution of expression values is statistically consistent with population level estimates over each subset of patients from the p-freedom estimation step. This is based on the assumption that the estimated distribution of cell-type-specific expression in each subset is robust to outliers.

Given that this solution is a function of the window size, which is an artificially defined parameter, we suspect that a consensus solution obtained from averaging an ensemble of solutions from different window sizes would be more robust and closer to the ground truth. Hence, in our ensemble sliding window deconvolution, we set up window sizes ranging from $s_1 = \frac{(1.5 \times number\ of\ cell\ types)}{0.8}$ times the number of cell types to $s_t = \max\left(4 \times \text{number of cell types}, \frac{\text{sample size}}{2}\right)$ and then perform the above sliding window deconvolution for each of these window sizes. Given multiple solutions for the cell-type-specific expression profile of each sample derived from multiple choices of sliding-window sizes ($s_1$ to $s_t$ ), their average is computed to obtain a single initial approximate solution to problem (1) (we refer to this as ensemble of window sizes). Steps 2.1-2.3 are repeated for the next gene until all the genes are done. Given a reimplementation of the CIBERSORTx sliding window algorithm (as function sliding_window()), the ensemble sliding window pseudo code is provided below:

**ensemble_sliding_window():**

**Input:**

$B$ = *bulk expression matrix*

$F$ = *cell fraction matrix*

$i$ = *gene index*

$n$ = *the number of samples*

$G^p$ = *expected cell-type-specific expression distribution over samples obtained from recursive splitting step*

$c$ = *the number of cell types;*

$m$ = *the number of genes;*

**Output:**

*Cell-type-specific expression matrix (3-dimension) for all samples:* $G$

***Function:***

$s_1 = \frac{(1.5 \times c)}{0.8}$;

$s_t = \max\left(4c,\ \text{round}\left(\frac{size}{2}\right)\right)$;

$G$ = *initialize matrix*($num \times size \times c, 0$);

*for iteration* $w$ *from* $s_1$ *to* $s_t$

*{*

$G = G +$ *sliding window*($B$, $F$, $G^p$, $w, i$)*;(See CIBERSORTx algorithm for more details)*

*}*

$G = G/((s\_t - s\_1 + 1))$;

*Return* $G$*;*

***End***

**3. Confidence ranking for the predictions following module 1**

We expect that genes that follow the modeling assumptions of module 1 are more likely to have their cell-type-specific expression levels predicted confidently. Hence, while executing module 1, we collect a series of features that could be useful in determining confidence level of expression predictions for each gene-cell-type pair. These are: p-value of t-test determining if a gene is weakly expressed in a cell type (obtained from completion of step 2.1), ratio of mean predicted expression levels and p-value of differential expression between subsets $h_1$ and $h_2$ (obtained from completion of steps 2.2 and 2.3), Spearman correlation between predicted cell-type-specific expression and bulk gene expression across samples, Spearman correlation between bulk expression and the cell fraction across samples etc. We then define a ranking $\Phi$ using this feature space such that genes achieving a high rank are on average ranked highly by each feature as follows:

$$\Phi^1\ (\text{gene } i, \text{celltype } k) = \left( \frac{\sum_{feature \in set} \text{rank}\big(feature(\text{gene } i, \text{celltype } k)\big)}{|set|} \right)$$

where $\Phi^1\ (\text{gene } i, \text{celltype } k)$ represents the prediction rank of gene $i$ in cell type $k$, $feature$ represents each feature we collected in the feature set, $|set|$ represents the number of features and $feature(\text{gene } j, \text{celltype } k)$ denotes each feature of gene $i$ in cell type $k$. The values taken by each feature are arranged so that for features representing p-values, lower the value higher the rank, but for features representing Spearman correlations, higher the value higher the rank.

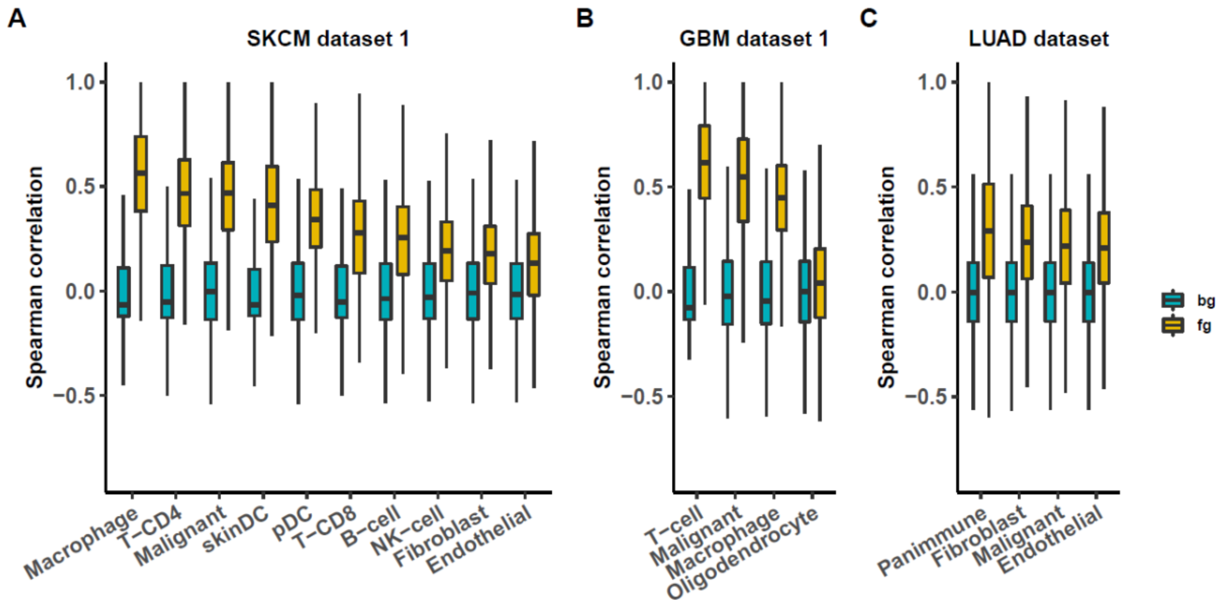

**Supplementary Figure S15: Gene-gene correlations among cell types. (A-C)** boxplots depicting the gene-gene expression correlation distributions among cell types in SKCM dataset 1, GBM dataset 1 and LUAD dataset respectively for 1000 randomly selected genes. In each of the three plots, corresponding random-permutation-based background controls are provided. The yellow box represents the correlation derived from the original datasets as the foreground (fg), while the green box represents that derived from the randomly permuted background control (bg). The y-axis denotes the Spearman correlation value and the x-axis denotes the cell type.

Additionally, it is well known that (the proteins encoded by) genes may interact with each other and behave collaboratively as complexes[29]; also, gene regulation is highly dependent on numerous regulatory elements including transcription factors[30,31]. When looking at single cell expression data from three independent single cell datasets, we indeed find that expression profiles of 1000 randomly selected genes within the same cell type are much more strongly correlated than expected by random chance (**Supplementary Fig. S15**). Hence, we reason that genes with correlated expression predictions for a given a cell type will have similar confidence levels. Therefore, ranking $\Phi$ is updated to $\Phi^1$ by accounting for these correlations as follows:

$$\Phi^1(\text{gene } i, \text{celltype } k) = \text{rank}_k(\max\{\Phi(\text{gene } j, \text{celltype } k): j \in Q\})$$

Here, $Q$ represents the set of genes whose predicted expression in cell type $k$ is strongly correlated with the predicted expression of gene $i$ in cell type $k$ (Spearman correlation ≥ 0.4). We now describe how the confidence ranking system takes this ranking $\Phi'$ and decides which genes need to be passed to module 2 for each cell type.

For each cell type $k$, we define two disjoint but non-exhaustive subsets: $\mathcal{H}_k$ and $\mathcal{L}_k$, which we call the “high” and “low”-confidence sets of cell type $k$. Genes belonging to the set $\mathcal{L}_k$ will be passed on to module 2. Let $m_k$ be the number of genes whose predicted expression distribution in the population is at least bimodal (i.e., fold change in expression between the subsets $h_1$ and $h_2$ > 1). Genes are then assigned to the high, low confidence set of each cell type by the confidence ranking system using the following rule:

$Add\ gene\ i\ to\ set$:

$$\mathcal{H}_k,\ \ if\ \Phi^1(gene\ i, celltype\ k) < \text{ round}\left(\frac{m_k^2}{2m}\right)$$

$$\mathcal{L}_k,\ \ if\ \Phi^1(gene\ i, celltype\ k) > m - \text{round}\left(\frac{m_k}{2} \times \left(1 - \frac{m_k}{m}\right)\right)$$

The results of the assignment are stored in a confidence matrix $C$ ($m$ genes $x$ $c$ cell types) encoding the high vs low confidence memberships of each gene in each cell type.

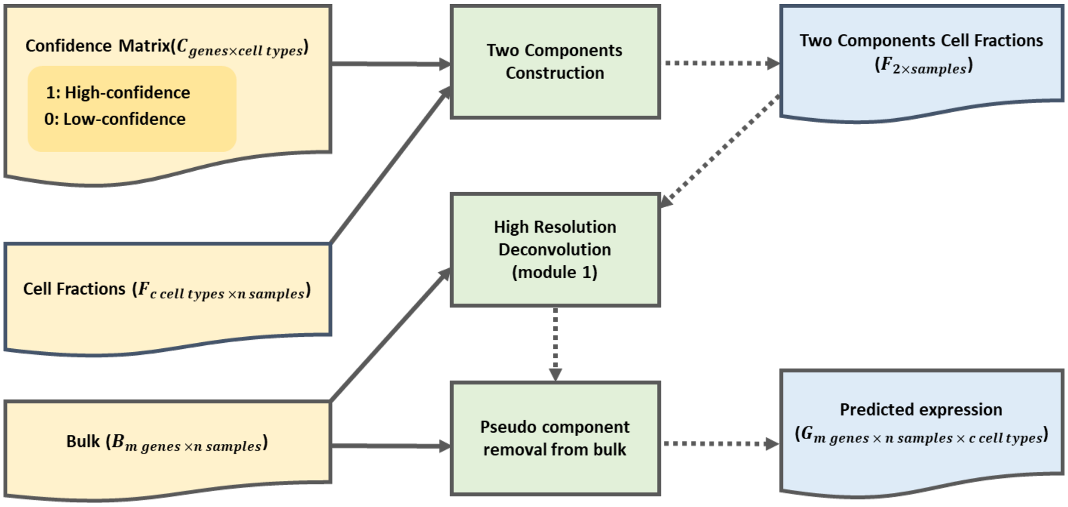

**Supplementary Figure S16: Hierarchical deconvolution.** Given the estimated cell fractions $F$ ($c$ cell types $\times$ $n$ samples), bulk and confidence levels estimated from module 1, for each cell type $k$ we merge all the other cell types as a pseudo component to construct a two-component model. Thereafter for each low-confidence gene $i$ in cell type $k$, we run module 1 to predict the expression in the pseudo component and finally remove the estimated expression of the pseudo component from the bulk to estimate the expression of the low-confidence gene $i$ in cell type $k$.

#### 4. Module 2 - Hierarchical deconvolution for low-confidence genes emerging from step 3

In this module, we simplify the general cell-type-mixture model described in module 1 to a 2-component mixture model (**Supplementary Fig. S16**). Specifically, for a gene $i$ in the low-confidence set of cell type $k$, its observed bulk expression level in a sample is modeled as a mixture of 2 components: the first component represents the cell type k and the second component represents a pseudo-cell-type that is a composite of all the cell types except $k^{th}$ cell type. We then re-run module 1 to predict individual specific expression of gene $i$ for the pseudo cell-type. Finally, the prediction for the pseudo cell-type is subtracted from the bulk to approximately re-estimate the individual specific expression of gene $i$ in cell type $k$. This is based on the assumption that the expression of the pseudo component might be better predicted than the expression of cell type $k$ using module 1, especially if cell type $k$ is not abundant or gene $i$ is weakly expressed in cell type $k$. The above steps are repeated for all the remaining genes in the low-confidence set of each cell type. The hierarchical deconvolution pseudo code is provided below:

**hierarchical_deconvolution():**

**Input:**

*$B$ = bulk expression matrix*

*$F$ = cell fraction matrix*

*$C$ = confidence level matrix (records low confidence genes in each cell type that need to be re-evaluated by module 2)*

*$G$ = predicted gene expression in each cell type and sample (Output of module 1)*

*$c$ = the number of cell types;*

*$m$ = the number of genes;*

**Output:**

*Updated predictions of gene expression in each cell type and sample in G*

***Function:***

```
for iteration k from 1 to c
{
    F" = [F[k,], 1 − F[k,]]
    for iteration d from 1 to m
    {
        if (C[i,k] == 0)
        {
            G^pseudo = High_resolution_deconvolution(B, F", i);
```

$$G[i,,k] = \frac{B[i,\ ] - F''[2,\ ] \times G^{pseudo}[i,\ ,2]}{F''[1,\ ]}$$

}

}

}

***End***

#### 5. Confidence ranking of predictions emerging from module 2

Following module 2, we re-rank all genes in the low confidence set $\mathcal{L}_k$ of each cell type by re-defining the ranking $\Phi^2$ as follows:

For gene $i \in \mathcal{L}_k$,

$$\Phi^2\ (gene\ i, celltype\ k) = rank_k(\frac{1}{|\mathcal{H}_k|} \sum_{j \in \mathcal{H}_k} \rho(\text{new predictions for gene } i, \text{predictions for gene } j))$$

Where $\rho$ represents the Spearman correlation, and $|\mathcal{H}_k|$ represents the number of genes in high confidence set $\mathcal{H}_k$. This is again based on observations of single cell expression data described above from which we deduce that genes with similar confidence levels are expected to have correlated predictions (**Supplementary Fig. S15**). We now describe how the confidence ranking system takes this new ranking of genes in the low confidence set of each cell type and decides which genes need to be upgraded to the high confidence set:

Let $|\mathcal{L}_k|$ be the number of genes in the low confidence set of cell type $k$, $N$ be the total number of genes and $CFM_k$ be the mean cell fraction of cell type $k$. The confidence ranking system upgrades the membership of genes from the low confidence set $\mathcal{L}_k$ to the high confidence set $\mathcal{H}_k$using the following rule.

$For\ gene\ i \in \mathcal{L}_k,$

$$\mathcal{H}_k \leftarrow \mathcal{H}_k \cup \{i\},\ \ if\ \Phi^2(gene\ i, celltype\ k) < \text{ round}\left(\frac{|\mathcal{L}_k|^2 * CFM_k}{2m}\right)$$

$$\mathcal{L}_k \leftarrow \mathcal{L}_k \ \{i\},\ \ if\ \Phi^2(gene\ i, celltype\ k) > m - \ \text{round}\left(\frac{|\mathcal{L}_k|^2}{2m}\right)$$

The results of the assignment are stored in the confidence matrix $C\ (m\ genes\ x\ c\ cell - types)$ encoding the high vs low confidence memberships of each gene in each cell type.

#### 6. Module 3 – Imputation-based deconvolution for low-confidence genes emerging from step 5

Module 3 operates on the assumption that the expression levels of two genes are supposed to be correlated in some cell types if we observe that their bulk expression is significantly correlated [16]. For a gene $i$ still in the low-confidence set of cell type $k$, the Spearman correlations between the bulk expression profile of gene $i$ and bulk expression profiles of genes in the high confidence set of cell-type $k$ are estimated. If the bulk expression profile of gene $i$ is highly correlated (Spearman correlation ≥ 0.5) with the bulk expression profiles of more than two genes in the high-confidence set of cell type $k$, then a lasso regression-based machine learning model is trained using bulk expression to impute individual specific expression of gene $i$ in cell type $k$ based on predicted expression profiles of high-confidence genes in cell type $k$ (**Supplementary Fig. S17**). The above steps are repeated for all the remaining genes in the low-confidence set of each cell type.

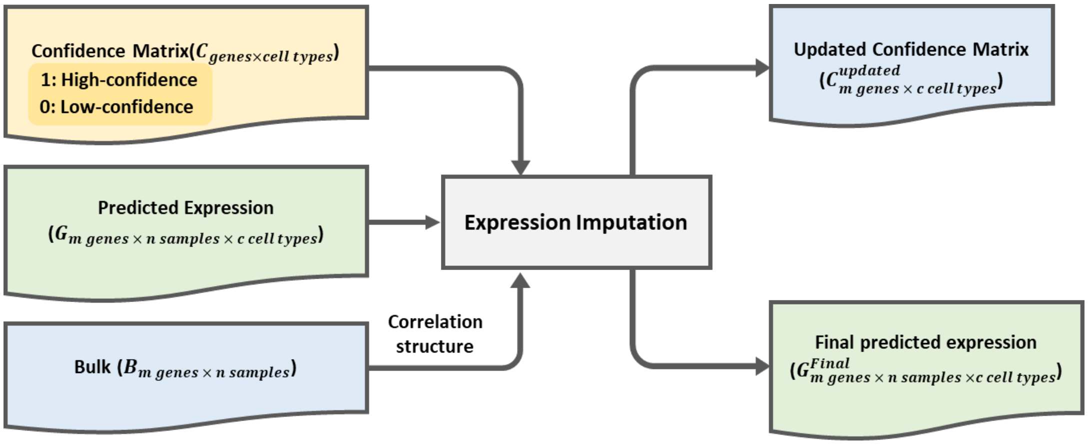

**Supplementary Figure S17: Imputation-based deconvolution.** Given the predicted cell-type-specific expression $G$ ($m\,\text{genes} \times n\,\text{samples} \times c\,\text{cell types}$), bulk and confidence levels estimated from module 1, in each cell type $k$ and for each low-confidence gene $i$, we compute the correlation between gene $i$ and each of other genes in bulk. If the number of genes which are highly correlated with gene $i$ is more than 2, we build up a machine learning model to predict the expression of gene $i$ in cell type k based on the expression of other high-confidence genes which are correlated with gene $i$. After imputation, both the predicted expression matrix $G$ and confidence matrix $C$ will be updated to record the final low/high confidence memberships of genes in each cell type.

**Imputation_based_deconvolution():**

**Input:**

$B$ = *bulk expression matrix*

$F$ = *cell fraction matrix*

$C$ = *confidence level matrix (records low confidence genes in each cell type that need to be re-evaluated by module 3)*

$G$ = *predicted gene expression in each cell type and sample (Output of module 2)*

$c$ = *the number of cell types;*

**Output:**

*Updated predictions of gene expression in each cell type and sample recorded in G*

***Function:***

*for iteration* $k$ *from 1 to* $c$

$\{$

$\mathcal{H}_k = gene\ j\ with\ \ C[j,k] == 1;$

$\mathcal{L}_k = gene\ j\ with\ \ C[j,k] == 0;$

$Conf_{high} = all\ genes \in\ \mathcal{H}_k;$

$Conf_{low} = all\ genes \in\ \mathcal{L}_k;$

$Corrs = pairwise\ Spearman\ correlation\ matrix\ (B[Conf_{low}, ] \times B[Conf_{high}, ]);$

*for each gene* $i \in \mathcal{L}_k$

{

*if* $(sum(Corrs[i,] \geq 0.5) \geq 2)$

{

*Train imputation model:* $B[i,] \sim f_{imp}(B[Conf_{high},]);$

*Impute* $G[i,,k] \leftarrow f_{imp}(G[Conf_{high},,k]);$

}

}

}

***End***

**7. Confidence ranking for predictions emerging from module 3**

Following module 3, we collect the following confidence ranking features for each gene $i$ in the low confidence set of cell type $k$: the correlations of predicted gene expression with bulk expression, the correlation between cell fractions and bulk expression, number of genes as features in the imputation model, average Spearman correlation between new predictions of gene $i$ and predictions of genes in the high confidence set of cell type $k$. We re-define a ranking $\Phi^3$ over all genes in the low confidence set of each cell type using this feature space such that genes achieving a high rank are on average ranked highly by each feature. Genes that are ranked among top 80% (an artificial cutoff) of all genes in the low confidence set of a cell type $k$ are now upgraded to the high confidence set of cell type $k$ by the confidence ranking system.

**8. Final output – confidence scores and cell-type-specific gene expression profiles of each sample**

To transform high- vs low-confidence set memberships of genes in each cell type (which were based on artificially defined rules/cut-offs for easy implementation of the greedy algorithm), into scores that are continuous in the range [0,1], the following final steps were taken: **(a)** The pair-wise correlations between the predicted expression profile of a gene $i$ in cell type $k$ and predicted expression profiles of genes belonging to the high-confidence set of cell type $k$ are averaged to generate a score for gene $i$; **(b)** the cell-type-specific expression predictions across the samples (columns) are randomly shuffled to generate a background and step *(a)* is repeated to estimate a background distribution of scores for each gene; **(c)** for each gene and each cell type, one can now determine an empirical p-value $pv$ based on this background distribution of scores. These p-values quantify the probability of a gene having high confidence predictions by random chance if its predictions are correlated with predictions of any other genes belonging to the high-confidence set of a cell type. The p-values are low for genes that are part of the high confidence set and high for genes part of low confidence set and intermediate for genes belonging to neither. Hence, we record $1 - pv$ as the final confidence score for each gene-cell-type pair. The final outputs of CODEFACS are the approximate solution for 3-dimensional matrix $G$ after execution of module 3 (imputation-based deconvolution) and confidence scores for each gene-cell-type pair.

**Comparing the performance of module 1 in CODEFACS to that of CIBERSORTx**

As we have mentioned above, in module 1 of CODEFACS we extended the high-resolution deconvolution approach presented previously in the CIBERSORTx algorithm, by generalizing their two-freedom estimation method to "$p$-freedom estimation" and modifying their sliding window method to ensemble sliding window deconvolution. By applying CODEFACS to the six benchmark datasets with sufficient sample size (SKCM dataset 2, SKCM dataset 3, SKCM dataset 4, GBM dataset 2, GBM dataset 3, GBM dataset 4; **Supplementary Fig. S18**, sample size = 100), we showed our recursive splitting ($p$-freedom estimation) and ensemble of window sizes methods substantially improved the prediction performance (in terms of the number of genes with Kendall scores ≥ 0.3). This improvement is seen especially in abundant cell types and is robust to artificial noise introduced in different pseudo-bulk expression datasets. (**Supplementary Fig. S18B-C** and **Supplementary Fig. S18E-F**).

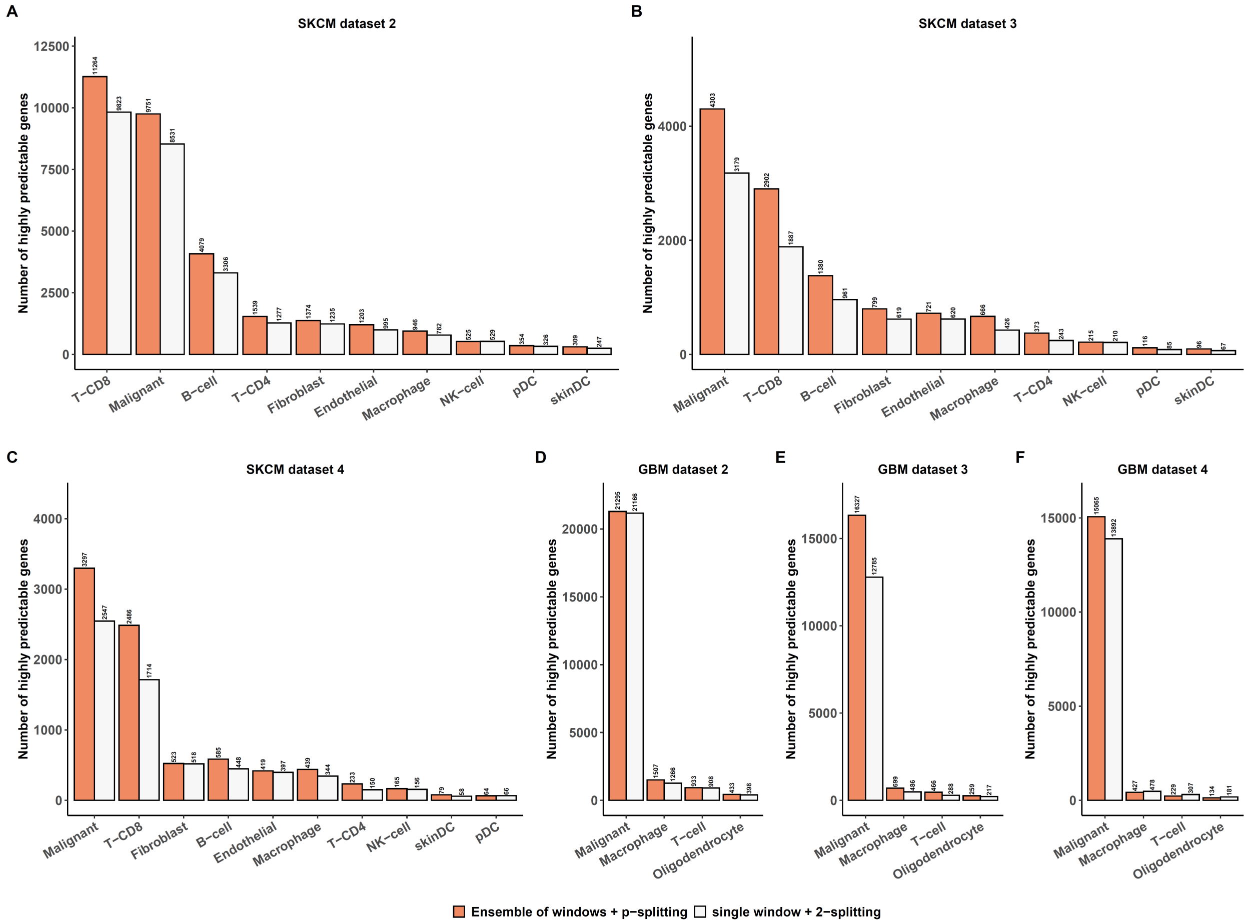

**Supplementary Figure S18: Performance comparison between two algorithm settings. (A-F)** bar plots depicting the number of highly predictable genes (Kendall correlation ≥ 0.3) using the two algorithmic approaches (ensemble of windows with p-splitting smoothing and single window with 2-splitting smoothing) in each cell type among six benchmark datasets with sufficient sample size (100). The orange bar represents the performance of ensemble of windows with p-splitting smoothing, while the white bar represents that of single window with 2-splitting smoothing. See **Supplementary Table 6** for more details on each benchmark dataset.

#### Demonstration of deconvolution performance gains as CODEFACS proceeds through each module

The CODEFACS algorithm relies on a series of heuristics that are expected to greedily predict closer to the ground truth as we sequentially execute each of its three modules. Here we show the performance of this approach in practice using the 15 benchmark datasets we generated by keeping track of the Kendal score distribution of high confidence genes at the end of each module (**Supplementary Fig. S19**). In general, for each cell type among the 15 benchmark datasets, we showed the overall predictions are improved across the three modules and module 2 achieves the most substantial improvement upon module 1.

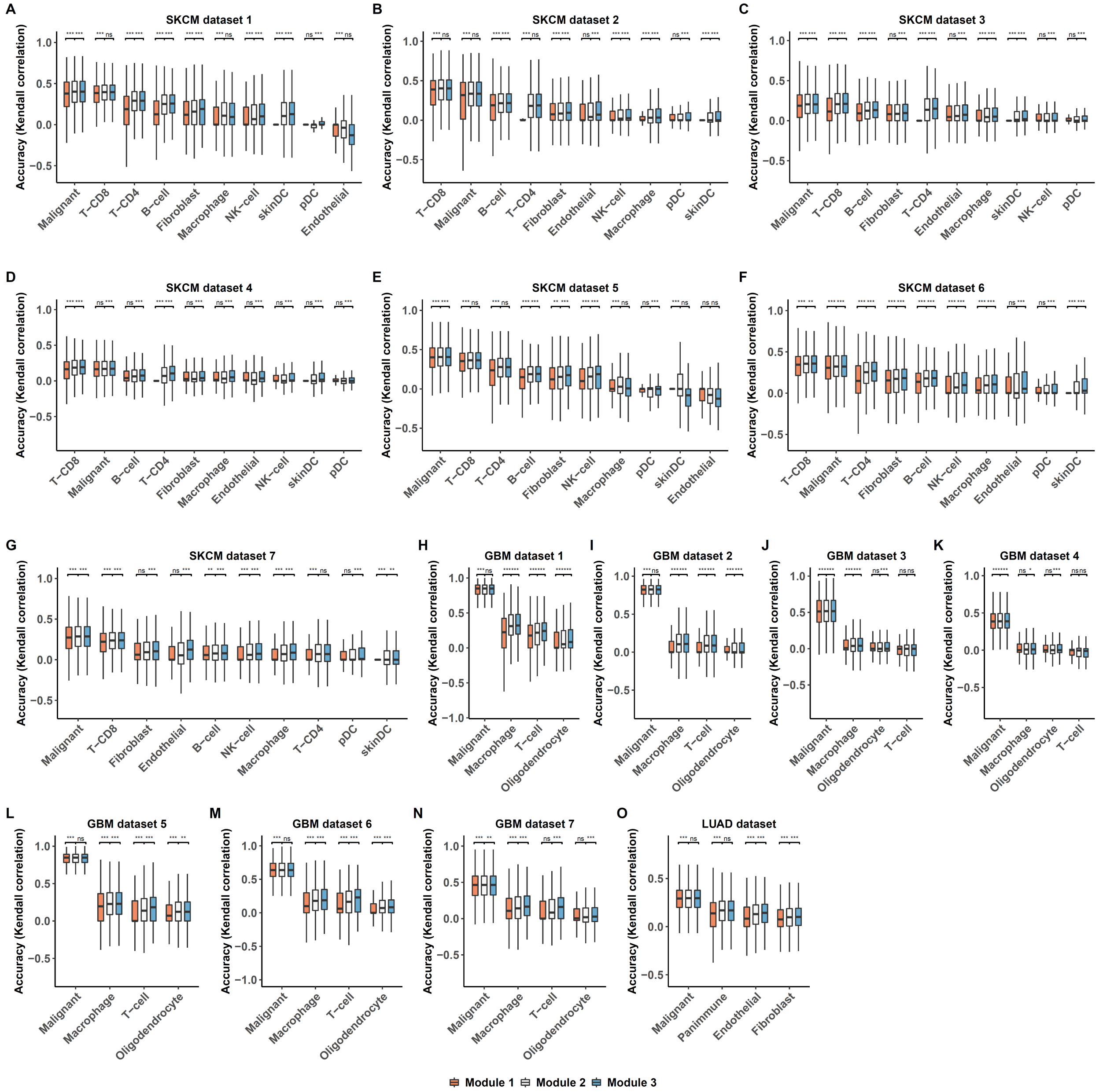

**Supplementary Figure S19: Prediction accuracy distributions across three algorithms steps in each cell type among the 15 benchmark datasets. (A-O)** boxplots depicting prediction accuracy distributions of high-confidence genes (based on CODEFACS) among cell types respectively in the 15 benchmark datasets. The orange bar represents the performance of module 1, the white bar represents that of module 2 and the blue one represents that of module 3. The y-axis indicates the prediction accuracy, and the x-axis indicates the cell types. See **Supplementary Table 6** for more details on each benchmark dataset.

### Supplementary Note 2: LIRICS (Ligand Receptor Interactions between Cell Subsets)

#### Curation of established ligand-receptor protein-protein interactions between cell types in the tissue microenvironment

Known protein-protein interactions between tumor, epithelial, immune and stromal cell types in the tissue microenvironment were manually curated from various resources. Specifically, interactions corresponding to cytokine/chemokine - cytokine/chemokine receptor interactions, ligand-receptor interactions involved in cell adhesion/leukocyte trans-endothelial migration, ligand-receptor interactions involving the TNF receptor superfamily and lastly, ligand receptor interactions involved in regulation of NK and T cell cytotoxicity were all merged into one Excel spreadsheet[32–38] (See **Supplementary Table 1**). In total, 369 putative ligand-receptor interactions were collected. This list primarily covers proteins that have well characterized immunological functions. Certain receptors are complexes encoded by more than one gene, such as TGF beta family receptors. They are documented as a list of genes separated by a ";". Furthermore, certain proteins serve as both ligands on some cell types and receptors on other cell types, such as HVEM (TNFRSF14).

#### Expected distribution of ligands and receptors across different cell types from prior knowledge

The database assigns a binary indicator (1/0), for each ligand/receptor, across the compendium of cell types indicating with 1 if the ligand/receptor can be produced by a cell type based on prior evidence of cell-surface protein expression or secretion (0 otherwise). This knowledge was extracted from Appendix II-IV of Janeway's Immunobiology 9th Edition Textbook[36] and systematically organized in **Supplementary Table 2**. The appendix, in addition, records ligands/receptors whose expected cell type specific distribution is less precisely defined. For instance, certain cytokines/chemokines are reported to be broadly produced by lymphocytes. Hence, without additional evidence, it is reasonable to expect that such ligands/receptors can also be produced in specific contexts by all cell types that are lymphocytes. We formalize this notion by defining a *functional equivalence class* for each cell type. For instance, the functional equivalence class for B cells is defined as: {lymphocytes, lymphoid cells, leukocytes, antigen presenting cells, nucleated cells, all cells}. A schema representing such relationships is stored in the third column in **Supplementary Table 3**. We then describe in a subsequent section how this prior knowledge can be used to systematically enumerate all ligands/receptors that can potentially be produced by a specific cell type of interest.

#### Annotation of functional effects of ligand-receptor interactions on participating cell types

Certain ligand-receptor interactions between immune cell types have an activating or inhibitory effect on the cell type expressing the receptor (also known as the target cell type) or in some cases both the ligand and receptor expressing cell types (regarded in literature as costimulatory). Discovery of such interactions resulted in the development of immune checkpoint blockade therapy such as anti-PD1 and anti-CTLA4 which has revolutionized cancer treatment. We systematically curated literature on all such interactions from[32,33,37–44] and classified them into two ontologies as follows:

- ***Activating/costimulatory*** encapsulates interactions with the following functional characteristics reported in literature: increased cytotoxicity, increased cytokine production, increased cell proliferation, increased cell survival, existence of immunoreceptor tyrosine-based activation motifs (ITAMs) in the cytoplasmic tail of the receptor.
- ***Inhibitory/checkpoint*** encapsulates the following functional characteristics reported in literature: decreased cytotoxicity, exhaustion, reduced cytokine production, decreased TCR signaling activity (for T cells), reduced cell proliferation, reduced cell survival, existence of Immunoreceptor tyrosine-based inhibitory motifs (ITIMs) in the cytoplasmic tail of the receptor.

In the database, interactions with conflicting effects reported on target cell types or for cases where the effect of the interaction depends on other factors were left unannotated. In addition to activating/inhibitory interactions, the database also annotates other interactions based on prior knowledge from Janeway's immunobiology 9$^{th}$ Edition Textbook[36].

- ***Pro-inflammatory*** interactions involving inflammation mediator cytokines such Interferon Gamma, TNF-alpha, IL1, IL12 and IL18
- ***Chemotaxis****:* cytokine/chemokine interactions involved in cell chemotaxis in regular or inflammatory conditions (responsible for lymphocyte infiltration)
- ***Cell-adhesion****:* interactions involved in cell adhesion/leukocyte trans-endothelial migration (responsible for extravasation from blood vessels to tissue)

##### LIRICS STEP 1: Querying all plausible ligand receptor interactions between any two cell types based on prior knowledge

In this step, we query all ligand receptor interactions that could potentially take place between two cell types A and B. The user can plug in the names of any two cell types whose names match with the names in the cell types compendium (**Supplementary Table 2**) and then LIRICS lists all ligand-receptor interactions that could potentially take place between cell types A and B. This list is determined by first finding which ligands/receptors can potentially be produced by each cell type (cell type A and B). To determine this, LIRICS first queries **Supplementary Table 2** which catalogues the expected distribution of ligands/receptors on different cell types. It then adds to this list any ligands/receptors that are expected to be found in the functional equivalence class of each cell type (see **Supplementary Table 3**). Given a set of all potential ligands and receptors on each cell type, LIRICS returns all known physical protein-protein interactions involving these ligands and receptors (from **Supplementary Table 1**).

##### LIRICS STEP 2: Identifying which plausible interactions are likely to occur (or "active") in each sample given deconvolved gene expression data from CODEFACS

Given a queried set of all plausible ligand receptor interactions between cell types (A,B,C,...): $\{(L_1^A, R_1^B), (L_2^A, R_2^C), \dots, (L_1^C, R_1^B), \dots, \}$, one can integrate this prior knowledge with deconvolved expression data from CODEFACS to infer which interactions are likely to occur in each sample as follows:

For any two cell types A and B with a plausible ligand-receptor interaction $(L)_z^A, R_z^B$, we define a binary indicator $Z_{(L)_z^A, R_z^B} \in \{0,1\}$, such that $I_{(L)_z^A, R_z^B} = 1$ if a physical interaction between $(L)_z^A, R_z^B$ is likely to take place in a sample, and has the value 0 otherwise. An interaction is considered likely to take place (synonym: "active") in a sample if the ligand $L_z^A$ is overexpressed in cell type A and receptor $R_z^B$ is over expressed in cell type B, in that sample. To determine if ligand $L_z^A$ and receptor $R_z^B$ are over-expressed in cell types A and B in a given sample, we use the median deconvolved expression of the ligand $L_z^A$ in cell type A over all input samples and likewise the median deconvolved expression of receptor $R_z^B$ in cell type B over all input samples as controls. Ligands such as cytokines and chemokines, can be secreted by cells and hence are not surface bound. However, we expect the levels of secreted cytokines/chemokines by a cell type to be proportional to their cell-type-specific gene expression. Furthermore, multiple genes are required to encode certain ligands/receptors, each gene being part of a specific subunit in the protein complex. For such ligands/receptors to be expressed, all genes required to build the ligand or receptor need to be expressed. Hence, their expression in a cell type is modeled as the minimum of the expression of individual genes constituting the ligand or receptor.

This approach has two key advantages, besides being biologically intuitive. First, the binary indicator is expected to be robust to noise in gene expression despite the varying levels of confidence in the predicted cell-type-specific gene expression from different datasets. This follows from the statistical properties of median-based filters in signal processing[45]. Second, it enables comparison of individual profiles independent of the dataset source due to their shared biological representation if the datasets being compared have expression measurements of the same genes. Thus, one can seamlessly pool multiple datasets together, augment sample size and increase statistical power.

##### Downstream enrichment analysis and visualization

Given this binarized representation, one can perform a Fisher's exact test to assess if any specific cell-cell interaction is more likely to occur in samples with a specific phenotype compared to a control group. This is quantified by computing the enrichment score, expressed as an odds ratio of the interaction in each phenotype of interest. A score around 1 indicates a neutral trend, a score >1 indicates enrichment of the interaction in the phenotype of interest and a score close to 0 indicates enrichment in the control group. Furthermore, the associated p-values of each test can be inspected post multiple hypothesis testing correction to identify any significant trends in the data. One can also plot the most significant trends occurring in a network where each edge represents a ligand-receptor interaction between two cell types and the thickness of the edge is proportional to the enrichment score of the interaction in a phenotype of interest. The circlize package in R is used to make these plots[46].

11. Bakker OB, Xu CJ, Koenen HJPM, Joosten LAB, Netea MG. Deconvolution of bulk blood

eQTL effects into immune cell subpopulations. *BMC Bioinformatics*. 2019;21(1):243. doi:10.1101/548669

22. Wang N, Hoffman EP, Chen L, et al. Mathematical modelling of transcriptional heterogeneity identifies novel markers and subpopulations in complex tissues. *Sci Rep*.

2016;6. doi:10.1038/srep18909

36. Murphy K, Weaver C. *Janeway's Immunbiology. 9th Edition*.; 2017.
